# Plasma Extracellular Vesicle Lipidomics Reveals an SMPD1-Driven Sphingomyelin Salvage Pathway Collapse in Parkinson’s Disease

**DOI:** 10.64898/2026.09.25.754317

**Authors:** Sanskriti Rai, Astha Chaubey, Sangita Paul, Sadaqa Zehra, Ananya, Roopa Rajan, Shashank Shekhar, Jagadeshwar Reddy T, Suman Jain, Sarika Gupta, Krishna K Inampudi, Neerja Rani, Anita Mahadevan, Gyan P Modi, Saroj Kumar

## Abstract

While lysosomal lipid dysregulation is implicated in Parkinson’s disease (PD), identifying the precise enzymatic drivers remains limited by the lack of subcellularly resolved metabolic mapping. Leveraging endosome-origin plasma extracellular vesicle (PsEV) lipidomics, we identified a profound bidirectional disruption of the ceramide-sphingomyelin (SM) axis in PD, driven by asymmetric upregulation of sphingomyelin phosphodiesterase 1 (SMPD1) and sphingomyelin synthase 1 (SGMS1). BODIPY-C5 SM tracking revealed that SMPD1 perturbation severely impairs lysosomal SM turnover. Crucially, this SMPD1-driven hydrolysis forces a salvage pathway collapse, bottlenecked by SPHK1/SGPL1 downregulation, which precipitates oxidative membrane damage and PINK1/Parkin-dependent mitophagy failure. We robustly validated this multi-pathway dysfunction across orthogonal *in vitro*, *in vivo* mouse, and postmortem human brain models. Furthermore, SMPD1/SGMS1 expression distinctly stratifies clinical phenotypes, marking the aggressive Postural Instability/Gait Difficulty (PIGD) subtype. Our findings establish SMPD1-mediated lysosomal impairment and ceramide-SM imbalance as a core biochemical axis in PD pathophysiology.

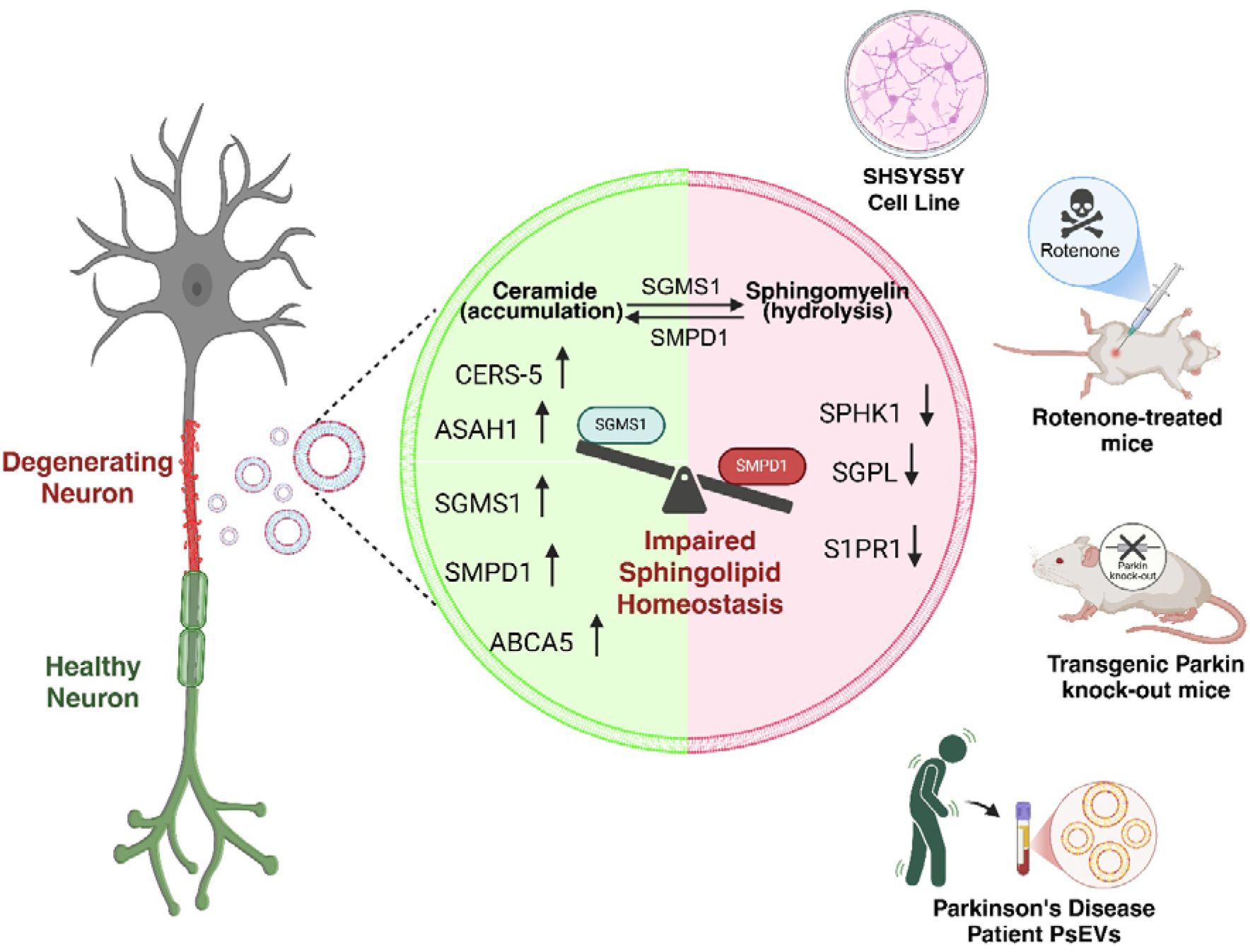

## Introduction

Idiopathic Parkinson’s disease (PD) ranks just behind Alzheimer’s disease among progressive neurodegenerative disorders, characterized mainly by bradykinesia, resting tremor, rigidity, and postural instability, which usually worsen as the disease progresses ^1^ ^2^. Its hallmark pathology includes degeneration of dopaminergic neurons in the substantia nigra pars compacta and the presence of intraneuronal Lewy body inclusions ^3^. Additionally, the pathological hallmark protein of Parkinson’s disease, α-synuclein, plays a crucial role in lipid membrane function ^4^.

Dysregulation of lipid dynamics within essential cellular pathways, including the endosome-lysosome system and synaptic signaling, contributes significantly to the manifestation of PD ^5^. Furthermore, lipid-modulating drugs, such as statins, are a topic of study for their protective effect against Parkinson’s disease ^6^. Numerous genetic predispositions associated with Parkinson’s disease include genes whose functions are related to lipid metabolism. Polymorphism within the genes GALC (which encodes galactosylceramidase) and ASAH1 (which encodes acid ceramidase), Sterol regulatory element binding transcription factor 1(SREBF1), another Parkinson’s disease risk gene that regulates sterol biosynthesis, critical for membrane integrity^7,8^, as well as mutations in GBA1, which encodes the lysosomal hydrolase glucocerebrosidase and represents the most common genetic risk factor for Parkinson’s disease ^9,10^. Dysregulation of lipid dynamics within essential cellular pathways, including the endosome-lysosome system, contributes significantly to the manifestation of PD. Among these pathways, sphingomyelin phosphodiesterase 1 (SMPD1),a critical lysosomal hydrolase responsible for degrading sphingomyelin into ceramide, has emerged as a key genetic and biochemical factor in PD susceptibility^11–13^.

Furthermore, meta-analyses examining targeted lipid investigations have revealed that elevated levels of total serum triacylglycerol and cholesterol exhibit a protective effect against the risk of Parkinson’s disease or are more prevalent in control groups as opposed to individuals with Parkinson’s disease ^14^. Serum lipids that differentiated Leucine-rich repeat kinase 2(LRRK2) mutation carriers and patients with Parkinson’s disease from controls were Ceramide species, Triacylglycerol, Sphingomyelin, Acylcarnitine, Phosphatidylcholine, and Lysophosphatidylethanolamine (LPE) and significant changes in Glycerolipids and Sphingolipids were associated with Parkinson’s disease severity ^15,16^. These findings imply an extensive dysfunction in lipid dynamics associated with Parkinson’s disease. Notwithstanding, a comprehensive characterisation of the overall lipidome specific to Parkinson’s disease is imperative for a balanced understanding of the involvement of lipids and sEVs association for newer insights into the disease diagnosis and understanding of the disease pathophysiology.

Studies from Indian cohorts have shown that Parkinson’s disease tends to begin at a younger age, with a mean age at onset of 53.64 ± 11.19 years, compared with the typically reported global range of 60-70 years, suggesting possible regional variation in disease presentation^17^. This observation is consistent with reports suggesting that genetic susceptibility, environmental exposures, and lifestyle factors may contribute to earlier disease onset in some populations^18^. Although pathogenic GBA variants are an established genetic risk factor for Parkinson’s disease, available Indian cohort studies indicate that their association with PD risk may be less consistent in the Indian population ^17,19,20^.

Despite this, current evidence remains limited by a reliance on bulk tissue or plasma lipidomics, which lack the subcellular spatial resolution required to pinpoint specific enzymatic failures. To overcome this spatial limitation, extracellular vesicles (EVs) have emerged as a powerful, minimally invasive tool. Because plasma-derived small EVs (PsEVs) originate predominantly from the endosomal-lysosomal pathway, their cargo provides a highly specific molecular window into intracellular lipid trafficking and lysosomal health. In addition, the available evidence remains fragmented, with scattered reports implicating individual pathway components as potential disease markers rather than elucidating an integrated mechanistic picture. There is also a relative lack of knockdown-based functional validation, and most studies have not examined these changes in parallel with related neurodegenerative disorders to delineate as PD-specific feature rather than more general consequence of neurodegeneration. Importantly, clinical correlation of the molecular findings has been largely absent, limiting their translational relevance.

In this study, we aim to provide a comprehensive, mechanistically integrated understanding of sphingomyelin pathway dysregulation in PD. By leveraging unbiased lipidomics on PsEVs from clinically stratified PD cohorts, we identified a distinct bidirectional disruption of the ceramide-sphingomyelin axis. To establish the functional consequences of this disruption, we performed targeted genetic knockdown of SMPD1 and utilized real-time metabolic tracking to map the resulting lysosomal failure. Finally, we validated this distinct molecular signature across orthogonal in vitro models, in vivo pharmacological and transgenic mouse models, and postmortem human brain tissue, demonstrating its direct correlation with aggressive clinical motor phenotypes

## Results

### PD is associated with increased PsEVs and altered neuronal proteins concentration

Purification and characterization of small extracellular vesicles (sEVs) were performed following the Minimal Information for Studies of Extracellular Vesicles (MISEV) guidelines ^21,22^ (**Figure 1A**). Study enrolled 90 individuals with clinically diagnosed PD alongside an age matched healthy control group (n=60), allowing case-control comparisons that are not confounded by age related changes in lipid metabolism. Within the PD cohort, patients were stratified both by motor phenotype and by disease stage. Based on established criteria, 48.9% of patients were classified as tremor dominant (TD), whereas 44.4% fell into the postural instability–gait difficulty (PIGD) subtype, providing sufficient representation of both major motor phenotypes for subtype specific analyses (**Figure 1B**). PD severity was graded using the Hoehn and Yahr (HY) scale, with participants spanning early to moderate disease stages; the majority of patients were at HY stage 2, consistent with a predominance of mild bilateral involvement without impairment of balance (**Figure 1C**). Demographic variables, including age and sex distribution, were comparable between PD (Age= 54.426 ±10.415) and control group (Age= 52.5 ±9.364), supporting robust interpretation of EV □ associated sphingolipid changes in relation to PD status rather than demographic bias. Transmission electron microscopy (TEM) revealed PsEVs as spherical vesicles with intact lipid bilayers (**Figure 1D**) and CD9-labelled EVs (**Figure 1E**). Nanoparticle tracking analysis (NTA) confirmed the size distribution (**Figure 1F)** and quantitatively demonstrated a significant elevation in PsEVs concentration in PD plasma samples relative to controls (p = <0.0001; **Figure 1G**). Notably, plasma from PD patients contained a higher concentration of PsEVs compared to both age-matched controls (**Figure 1H)**. Western blot analysis validated the identity of isolated PsEVs through the detection of classical EV markers CD9. CD63, and Alix B (**Figure 1I**) (**Supplementary Figure S1A-C**). Densitometry analyses for CD9 (p= 0.02; **Figure 1J**), CD63 (p= 0.04; **Figure 1K**) and Alix (p= 0.04; **Figure 1L**) showed significant increase in PD patient. EV exclusion marker Calnexin B (**Figure 1M, Supplementary Figure S1D**) and protein co-precipate Apolipoprotein-B (**Figure 1N, Supplementary Figure S1E**) were also measured.

**Figure 1:**
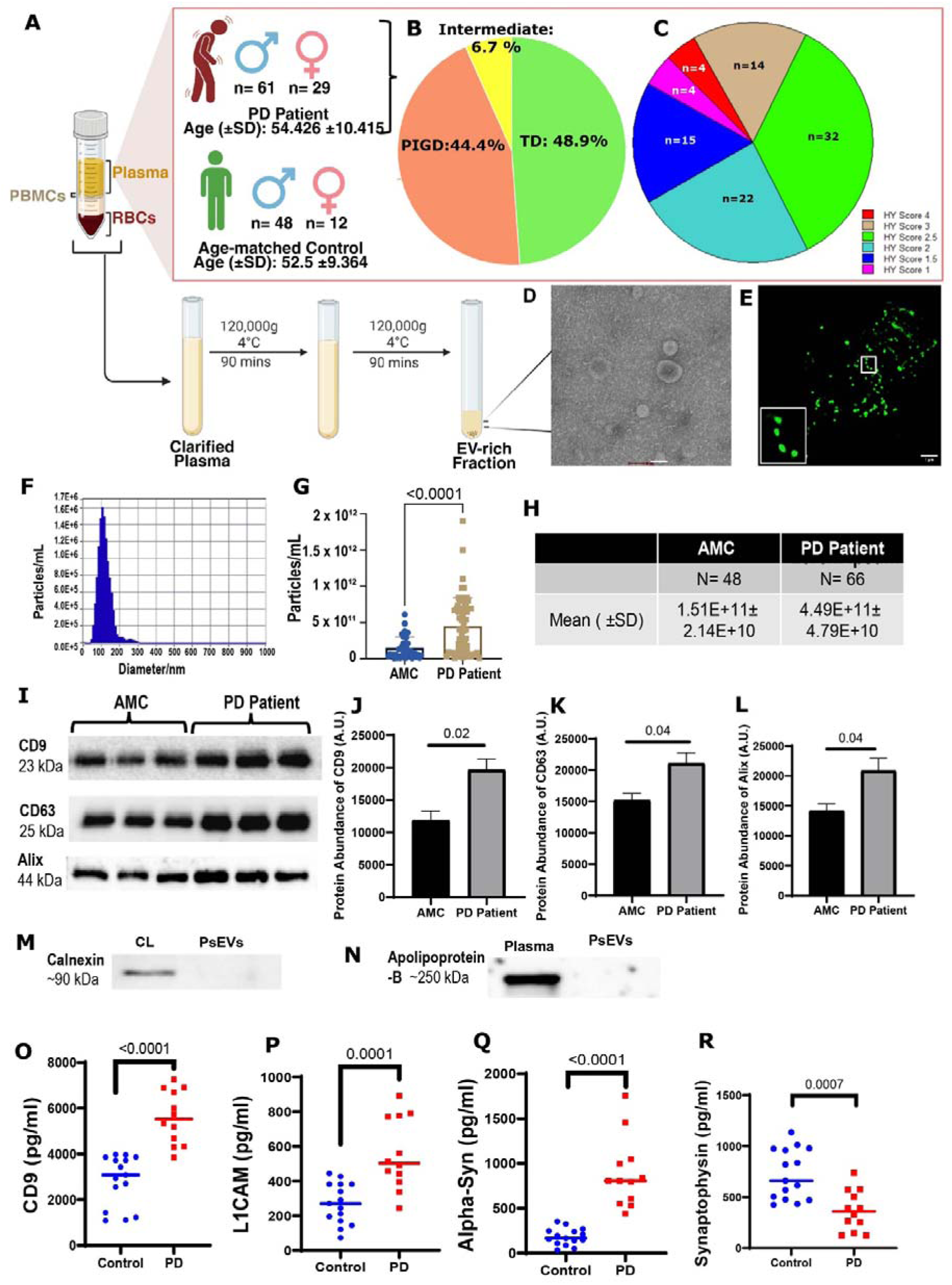
Isolation and characterization of plasma small extracellular vesicles (PsEVs) and profiling of EV-associated proteins in PD. (A) Schematic workflow of EV isolation. (B) Demographic characteristics of the study cohort and distribution of clinical subtypes. (C) Distribution of Parkinson’s disease (PD) patients according to Hoehn and Yahr (HY) stage. (D) Representative transmission electron microscopy images showing the morphology of isolated plasma sEVs (Scale bar= 100 nm). (E) Immunofluorescence-based detection of CD9-labeled EVs (Scale bar=1 μm). (F) Nanoparticle tracking analysis (NTA) showing the size distribution of PsEVs. (G, H) Particle concentration in control and PD samples. (I) Western blot analysis of the canonical EV markers CD9, CD63, and Alix. (J-L) Densitometric quantification of CD9, CD63, and Alix. (M) Calnexin, used as an exclusion marker. (N) Apolipoprotein B, a co-isolated protein marker. (O-R) ELISA-based quantification of CD9, L1CAM, α-synuclein, and Synaptophysin. Data are presented as Mean ± S.E.M. Statistical significance was assessed using Unpaired t-test, and p values are indicated.

Additionally, quantitative ELISA measurements of EV-associated proteins revealed a distinct PD-associated signature. EV marker CD9 levels were significantly increased in EVs from PD patients compared with controls (p<0.0001; **Figure 1O**), as were neuronal marker L1CAM (p = 0.0001; **Figure 1P**) and α synuclein (p < 0.0001; **Figure 1Q**), indicating an overall enrichment of neuronal and α synuclein–bearing vesicles in the PD group. In contrast, Synaptophysin levels were significantly reduced in PD-derived EVs (p = 0.0007; **Figure 1R**). Together, these data show that PD plasma EVs carry higher amounts of canonical tetraspanin (CD9), neuronal adhesion marker (L1CAM), and α synuclein, but lower Synaptophysin. Synaptophysin is a marker of synaptic vesicle integrity, its reduction in circulating EVs reflects the synaptic degeneration characteristic of PD, while the simultaneous enrichment of α-synuclein and L1CAM highlights the active vesicular packaging of pathological stress markers. This EV protein profile is consistent with the disease context, in which α synuclein-rich vesicles and altered synaptic integrity are expected to accompany neurodegeneration and changes in neuronal connectivity.

### Lipidomic profiling of PsEVs reveals elevated ceramide and disrupted sphingolipid metabolism in Parkinson’s disease

To elucidate the lipidomic profile of Plasma small extracellular vesicles (PsEVs), we performed comprehensive lipidomic analysis on PsEVs isolated from Parkinson’s disease (PD) patients and age-matched healthy controls (AMC). Lipid extraction was done using the modified Folch method, followed by shotgun lipidomics. Data normalization was done using internal standards, enabling absolute quantification of over 500 distinct lipid species spanning more than ten lipid classes. Principal component analysis (PCA) demonstrated clear separation between PD and AMC samples, with robust clustering within each group on the two-dimensional score plot, indicating high technical and biological reproducibility (**Figure 2A**). The PCA further confirmed distinct segregation of PD and AMC groups, reflecting pronounced differences in overall lipid composition. Analysis of the PsEV lipidome revealed that, when comparing disease and control conditions, 6.2% of lipid species were upregulated, 19.8% were downregulated, and 74% remained unchanged (**Figure 2B**). Lipidomic data were normalized to median values, log-transformed (base 10), and auto-scaled to facilitate comparison of lipid profiles between groups. Univariate statistical analyses were performed to further assess PsEV lipidome differences. We evaluated enrichment of lipid classes within PsEVs by calculating the percentage of each class in the total lipid content. As depicted in (**Figure 2C**), ceramide emerged as a major differentially expressed lipid class in the disease state. Dysregulation was also observed among several phospholipid classes, including phosphatidylethanolamine (PE), phosphatidylinositol (PI), phosphatidylserine (PS), phosphatidylcholine (PC), phosphatidic acid (PA), and phosphatidylglycerol (PG). Quantitative analysis revealed that ceramide levels were significantly elevated in PD samples compared to controls (p□ = □0.001; **Figure 2D**), while Sphingomyelin levels were significantly reduced in PD (p=0.0018; **Figure 2E**).

**Figure 2.**
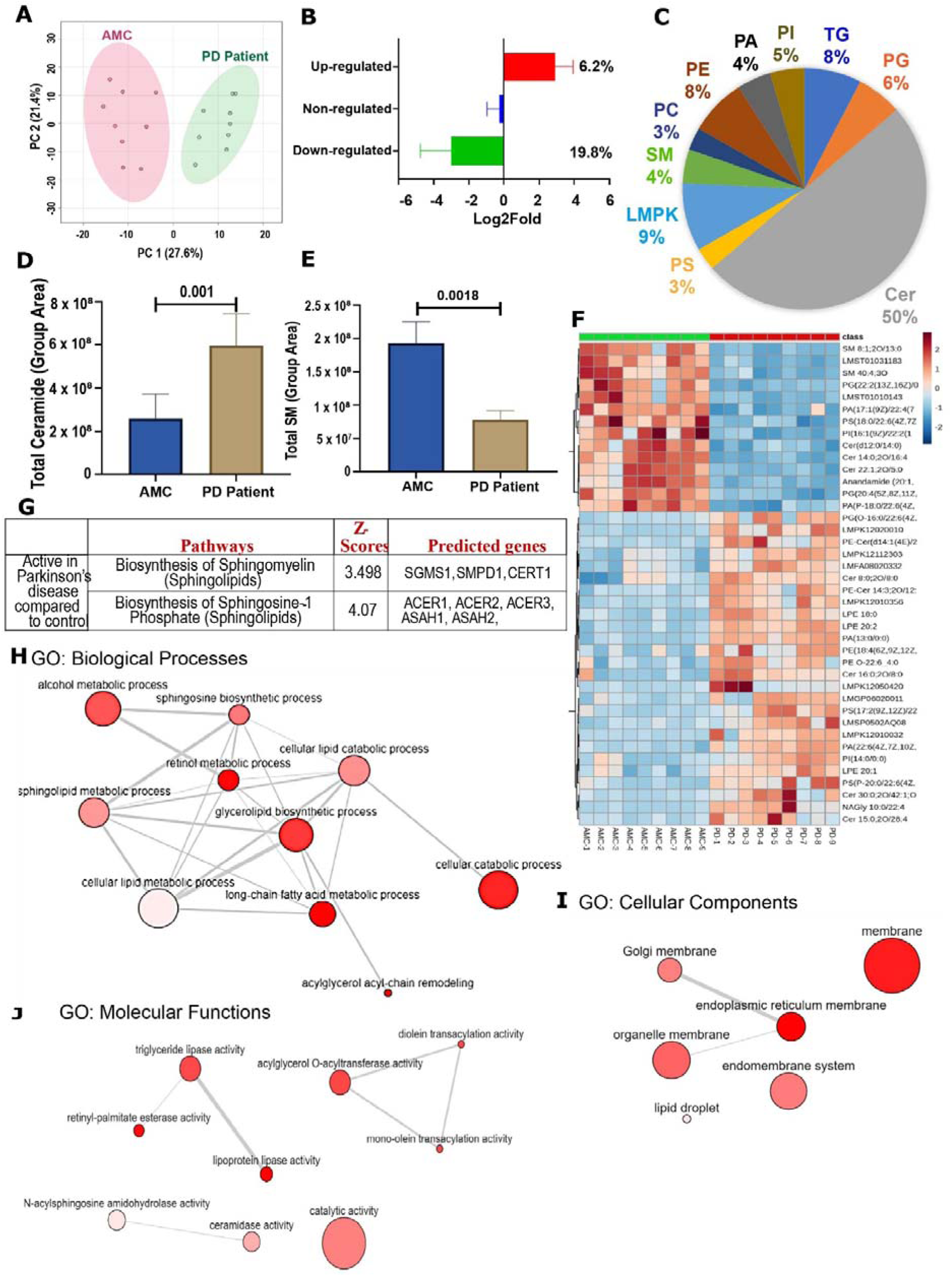
PsEV-derived lipidome and coverage of PD and AM-C. (A) Principal component analysis showing distinct separation between PD and control groups. (B) Total percentage of dysregulated lipids observed in Parkinson’s disease. (C) Pie chart showing the top 10 dysregulated lipid classes. (D) Total ceramide levels. (E) Total sphingomyelin levels. (F) Heat map showing the top 25 dysregulated lipid classes. (G) Table showing the sphingolipid metabolism pathway and predicted genes involved with a Z value > 1.96. (H-J) Gene ontology (GO) enrichment analysis of the identified genes, including biological processes, molecular functions, and cellular components.

Fold-change analysis identified 195 lipid species that were significantly altered (p□<□0.05) between PD patients and AMC (**Supplementary Figure S2A**). A heatmap of lipid-class composition (**Figure□2F**) highlights the diversity and differential abundance of PsEV-associated lipids between groups. Distinct expression patterns were evident, with notable alterations in polyketides (LMPK) and triglycerides when comparing PD and AMC profiles. Partial least squares-discriminant analysis (PLS-DA) was used to assess group separation and identify lipid species that contributed most strongly to discrimination between groups. (**Supplementary Figure S2B**). PLS-DA score plot demonstrated clear segregation between PD and AMC samples, with confidence ellipses indicating strong group clustering. Lipids with the highest contribution to group separation are shown in the VIP score plot (top 20 VIP scores; **(Supplementary Figure S2C**). Univariate statistical analyses were applied to further evaluate these differences. As illustrated in the volcano plot (**Supplementary Figure S2D**), each point represents a unique lipid species, with the x-axis indicating fold change and the y-axis corresponding to statistical significance (p□≤□0.05). Among these, 39 lipids were upregulated and 34 were downregulated in PD relative to controls

Differentially expressed lipids identified by LC-MS were analyzed using BioPan to assess known and predicted interactions, pathway activity, and associated genes. In PD compared with AMC, biosynthesis of sphingomyelin (sphingolipid metabolism) and biosynthesis of sphingosine□1□phosphate (sphingolipid metabolism) were predicted to be activated (**Figure□2G**). In contrast, pathways associated with Triglycerol catabolism and biosynthesis (Triglycerol metabolism), as well as biosynthesis of Phosphatidylethanolamine (PE), Phosphatidylcholine (PC), and Phosphatidylserine (PS) (Glycerolipid and Glycerophospholipid metabolism), were predicted to be suppressed in PD. BioPan further identified eight predicted genes with Z□scores ≥□1.68, the minimum accepted threshold (**Table□3**). To investigate potential interactions, these gene products were subjected to STRING analysis. The resulting protein–protein interaction network comprised 19 nodes and 41 edges, with highly significant enrichment (p□<□1.0□×□10^-^¹; (**Supplementary Figure S2E**). Gene Ontology (GO) (**Figure 2H-J**), and Kyoto Encyclopedia of Genes and Genomes (KEGG) (**Supplementary Figure S2F**). Enrichment analyses identified lipid metabolic processes as the dominant enriched category. In particular, sphingolipid metabolism and sphingolipid signaling pathways were significantly altered in PD compared with AMC.

### Alterations in Enzymes Regulating Ceramide and Sphingomyelin Metabolism in Parkinson’s Disease

To explore potential mechanisms underlying the dysregulated ceramide and sphingomyelin (SM) levels observed in PD PsEVs, we examined the expression of key enzymes involved in both the de novo and salvage pathways of ceramide synthesis (**Figure□3A**). Ceramide synthase□5 (CERS5) was significantly upregulated in the PD group (**Figure 3B**), whereas no difference was observed in the expression of serine palmitoyl transferase (SPTLC) and Ceramide synthase□4 (CERS4) (**Supplementary Figure□S3A-B**). We next assessed enzymes regulating sphingolipid synthesis and metabolism. Sphingomyelin synthase□1 (SGMS1), which catalyzes SM synthesis from ceramide, and sphingomyelin phosphodiesterase□1 (SMPD1), which mediates SM degradation, were both upregulated in PD relative to AMC (**Figure□3C-D**). To further understand the lysosomal handling of these lipids, we evaluated ABCA5, a critical lysosomal transporter responsible for sphingomyelin efflux. Expression of ABCA5 was elevated in PD (**Figure 3E**), likely representing a compensatory response to the altered intralysosomal sphingomyelin levels.

**Figure 3:**
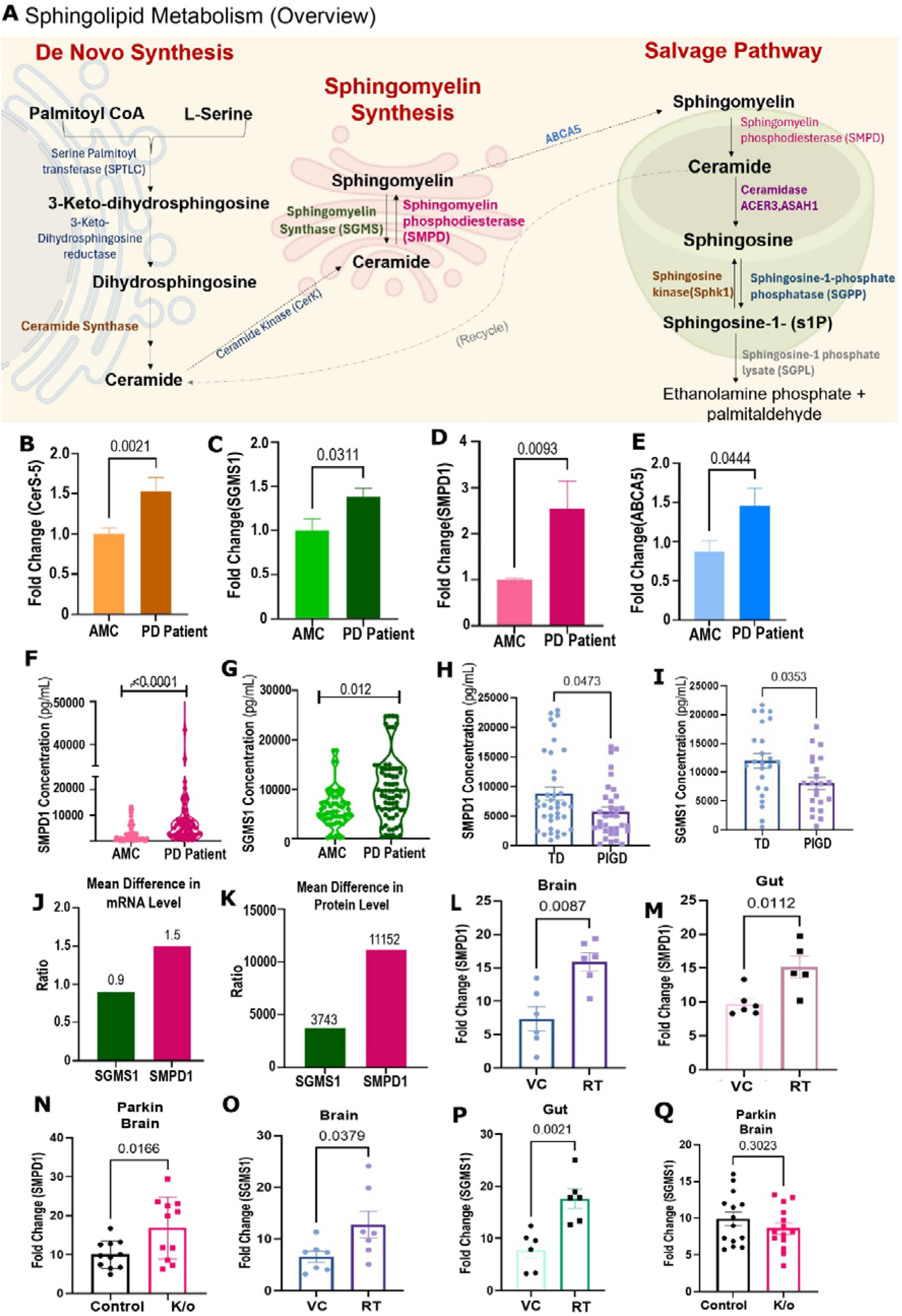
Sphingolipid metabolism and differential expression of key pathway components in PsEVs and mouse models. (A) Schematic representation of sphingolipid metabolism. (B–E) mRNA expression profiling of ceramide synthase-5 (CerS5), sphingomyelin synthase-1 (SGMS1), sphingomyelin phosphodiesterase 1 (SMPD1), and ABCA5. (F, G) Protein levels of SMPD1 and SGMS1 in PsEVs derived from PD patients and control individuals. (H, I) Protein expression of SMPD1 and SGMS1 across PD clinical subtypes. (J, K) Mean differences in mRNA and protein levels. (L-N) mRNA expression of SMPD1 in rotenone-treated mouse brain and gut, as well as in parkin knock-out brain. (O-Q) mRNA expression of SGMS1 in rotenone-treated mouse brain and gut, and in parkin knock-out brain. (AMC= Age-matched control, PD= Parkinson’s disease, TD= Tremor-dominant, PIGD= Postural instability and gait disability, VC= Vehicle Control; RT= Rotenone-treated mice). Data are presented as Mean ± S.E.M. Statistical significance was determined using Unpaired t-test, and p values are indicated

To investigate sphingolipid metabolism in PD, we quantified protein levels of sphingomyelin synthase□1 (SGMS1) and sphingomyelin phosphodiesterase□1 (SMPD1) in PsEVs from PD patients and AMC. SMPD1 protein level was significantly higher in the PD group (p<0.0001; **Figure□3F**), and SGMS1 protein levels were also elevated in PD (p□=□0.012; **Figure□3G**). Notably, significant differences in the expression of both enzymes between Tremor dominant (TD) and Postural instability and gait difficulty (PIGD) groups. SMPD1 protein levels were significantly reduced in the PIGD group compared to the TD group (p□=□0.0473; **Figure 3H**), as well as SGMS1 protein levels (p□=□0.0353; **Figure 3I**). Consistently, the protein levels of both SGMS1 and SMPD1 were markedly downregulated in the PIGD subtype, underscoring a distinctive sphingolipid enzyme profile associated with this clinically relevant motor phenotype.

Comparison of mRNA expression between the two enzymes revealed a 1.5□fold higher level of SMPD1 relative to SGMS1(**Figure□3J**). Notably, at the protein level, SMPD1 upregulation was more pronounced than that of SGMS1 (**Figure□3K).**

To determine whether this SMPD1/SGMS1 dysregulation is a unique feature of PD or a general consequence of neurodegeneration, we evaluated the expression of these enzymes in multiple sclerosis (MS) and Alzheimer’s disease (AD). While SGMS1 (**Supplementary Figure S3C**) and SMPD1 expression was unchanged in MS (**Supplementary Figure S3D)**, SMPD1 was significantly elevated in AD (p=0.0022; **Supplementary Figure S3E)**. However, the overall sphingolipid enzymatic axis exhibited a distinctly disease-specific topology. In PD, SMPD1 elevation is accompanied by a coordinated, albeit asymmetric, upregulation of SGMS1. In contrast, AD samples exhibited a unidirectional dysregulation characterized by elevated SMPD1 coupled with a significant decrease in SGMS1 (p=0.0037; **Supplementary Figure S3F)**. These findings demonstrate that while isolated lysosomal hydrolase hyperactivity (SMPD1) may overlap with AD, the bidirectional, synergistic disruption of the ceramide-SM axis, where asymmetric enzyme upregulation actively tilts homeostasis, thus making a specific molecular signature of PD

To evaluate whether the sphingomyelin pathway dysregulation observed in circulating human PsEVs reflects localized central nervous system pathology or a broader systemic phenomenon, we analyzed tissue from *in vivo* PD models. Given the emerging prominence of the gut-brain axis in PD pathophysiology, we assessed sphingomyelin pathway members not only in the brain but also in the gut tissues of rotenone treated mice (**Supplementary Figure S4A-H)**, alongside brain lysates from parkin knockout mice (**Supplementary Figure S5A-J)**. SMPD1 expression was significantly increased in both the brain (**Figure 3L**), and the gut of rotenone treated animals (**Figure 3M**) which was also elevated in the parkin knockout brain (**Figure 3N**). SGMS1 was upregulated in brain (**Figure 3O**) and gut (**Figure 3P**) of rotenone treated mice. We also assessed SGMS1 level in parkin knockout brain (**Figure 3Q**). Furthermore, correlation analysis revealed that PsEVs concentration was positively associated with SMPD1 (r= 0.4239; p<0.0001), SGMS1 (r= 0.33214; p=0.008), and α synuclein (r = 0.5158; p<0.0001), indicating that higher vesicle abundance accompanies increased levels of these sphingomyelin pathway and PD related proteins (**Supplementary Figure S3O-Q**). A similar positive correlation was observed between SMPD1 and SGMS1 expression (r= 0.44; p<0.0001) (**Supplementary Figure S3R**), supporting coordinated modulation of these enzymes within the same samples. No significant differences in SMPD1 or SGMS1 levels were detected when stratifying patients by sex or by familial versus sporadic PD status (**Supplementary Figure S3S-V**), suggesting that these associations are not driven by basic demographic or inheritance patterns.

### Salvage Pathway Dysregulation and Ceramide Accumulation in Human and Mouse PD Samples

To further investigate the enzymatic drivers of the observed ceramide-sphingomyelin imbalance, we examined the expression of key enzymes regulating salvage pathway of sphingolipid metabolism (**Figure□4A**). Acid ceramidase (ASAH1), responsible for ceramide degradation, was also upregulated in PD (**Figure□4B**) whereas, sphingosine kinase 1 (SPHK1), which converts ceramide to sphingosine, was downregulated (**Figure□4C**). Concomitantly, S1PR1 (**Figure□4D**) and SGPL1 (**Figure□4E**) were also decreased in PD, indicating compromised sphingosine-1-phosphate signaling and catabolism. Parallel analyses in mouse models revealed SGPL1 mRNA was significantly decreased in the brains of both rotenone treated (**Figure 4F**) and parkin knockout mice (**Figure 4G**). S1PR1 expression in the brain was likewise reduced in both models (**Figure 4H, I**). Lipidomic profiling showed a decrease in total sphingomyelin accompanied by selective enrichment of ceramide species, significantly Cer 16:0, and Cer 20:1 (**Figure 4J**). To explore whether these lipid changes were associated with altered mitochondrial quality control as reported by studies ^23^, we assessed PINK1 expression in the SMPD1 knockdown model and found it to be reduced relative to untreated controls (p = 0.02; **Figure 4K,L**; **Supplementary Figure S6A-C**). Parkin levels, however, were not significantly changed under the same conditions (**Supplementary Figure S6D-F)** compared with untreated controls. Collectively, the results indicate engagement of the sphingomyelin salvage pathway, and accumulation of bioactive ceramide species, due to concomitant attenuation of sphingosine-1-phosphate signaling., which is consistent with a shift in ceramide-sphingomyelin homeostasis.

**Figure 4:**
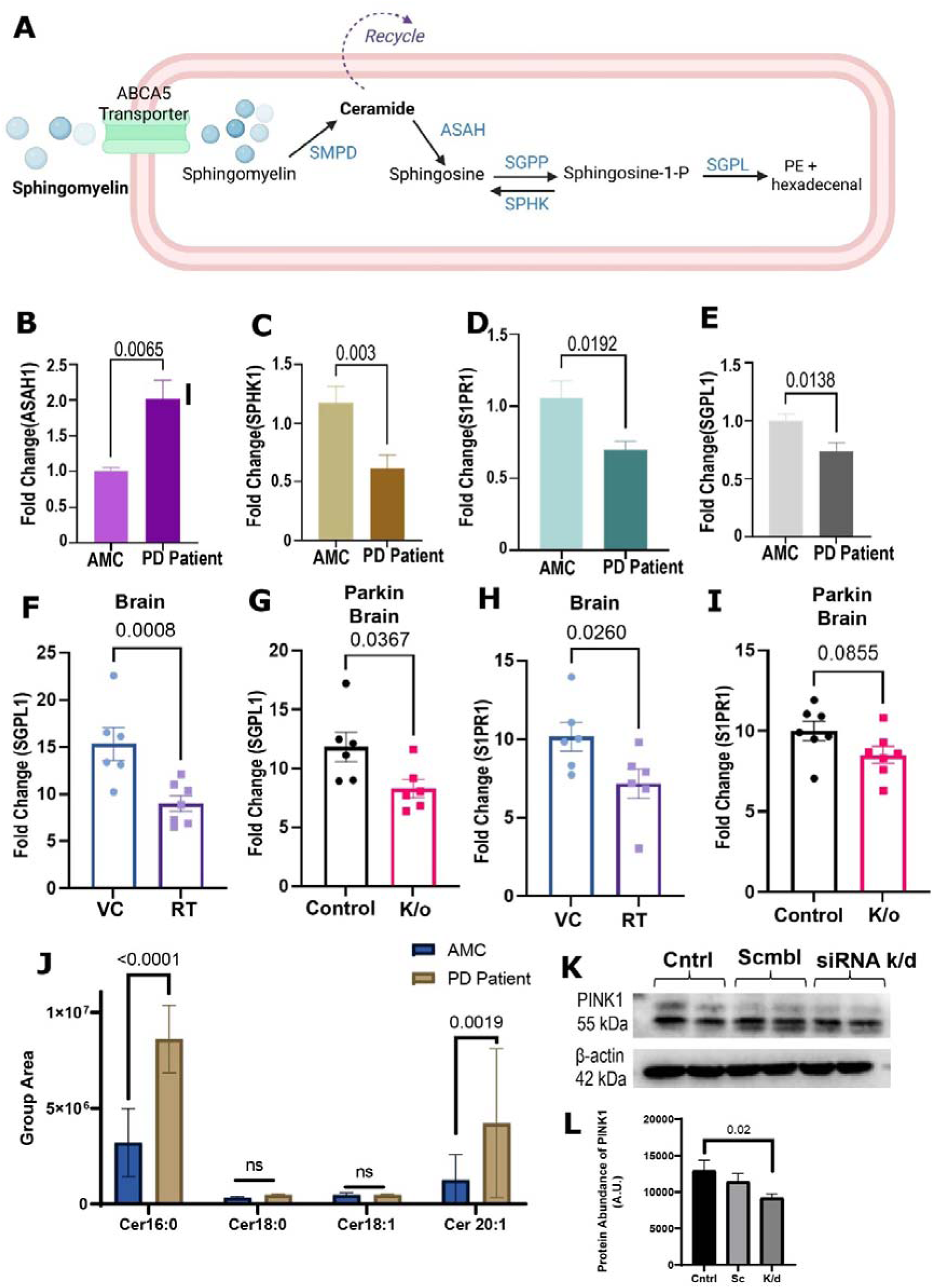
Sphingomyelin salvage pathway and associated molecular changes in PD patient-derived samples and mouse models. (A) Schematic representation of the sphingomyelin salvage pathway. (B-E) mRNA expression profiling in PD patient- and control-derived samples for ASAH1, SPHK1, S1PR1, and SGPL1. (F, G) mRNA expression of SGPL1 in rotenone-treated mouse brain and parkin knock-out brain. (H, I) S1PR1 levels in rotenone-treated mouse brain and parkin knock-out brain. (J) Profile of different ceramide species. Data are presented as Mean ± S.E.M. Statistical significance was determined using Unpaired t-test with p values indicated. (K) Western blot of PINK1 (L) Densitometry analysis of western blot. (AMC= Age-matched control, PD= Parkinson’s disease, VC= Vehicle Control; RT= Rotenone-treated mice, k/o= Parkin knockout, Cntrl= Control, Sc= Scramble, k/d= siRNA knock-down). Data are represented as Mean ±SEM, p value as indicated.

### SMPD1 knockdown impairs sphingomyelin catabolism and promotes lipid peroxidation in neuronal models

To evaluate the downstream functional consequences of SMPD1 perturbation, SMPD1 knockdown was performed in the SH-SY5Y cell line using targeted siRNA (**Figure 5A**). Both SMPD1 protein (**Figure 5B, Supplementary Figure 7A-C**) and mRNA levels were significantly decreased at 48 h compared to cells treated (p=0.0286) with a non-targeting Scramble (SC) control RNA (p=0.0383; **Figure 5C**). Next, we monitored oxidative membrane damage using the ratiometric lipid peroxidation probe BODIPY 581/591 C11. This probe undergoes a spectral shift from red to green fluorescence upon active lipid peroxidation. Compared to the Scramble (SC) control cells, SMPD1-knockdown cells exhibited a significant increase in green fluorescence (p=0.01; **Figure 5D**). This demonstrates that impaired sphingomyelin catabolism directly promotes elevated oxidative membrane stress. Observed spectral shift reflects oxidation of the diene bond in BODIPY, leading to loss of conjugation between the phenyl moiety and the boron dipyromethene difluoride core; the oxidized probe, in this altered form, exhibits green fluorescence. The increase in green fluorescence was significant after 30 minutes of incubation. No change in BODIPY green fluorescence was observed in untreated control wells (**Figure 5E**).

**Figure 5:**
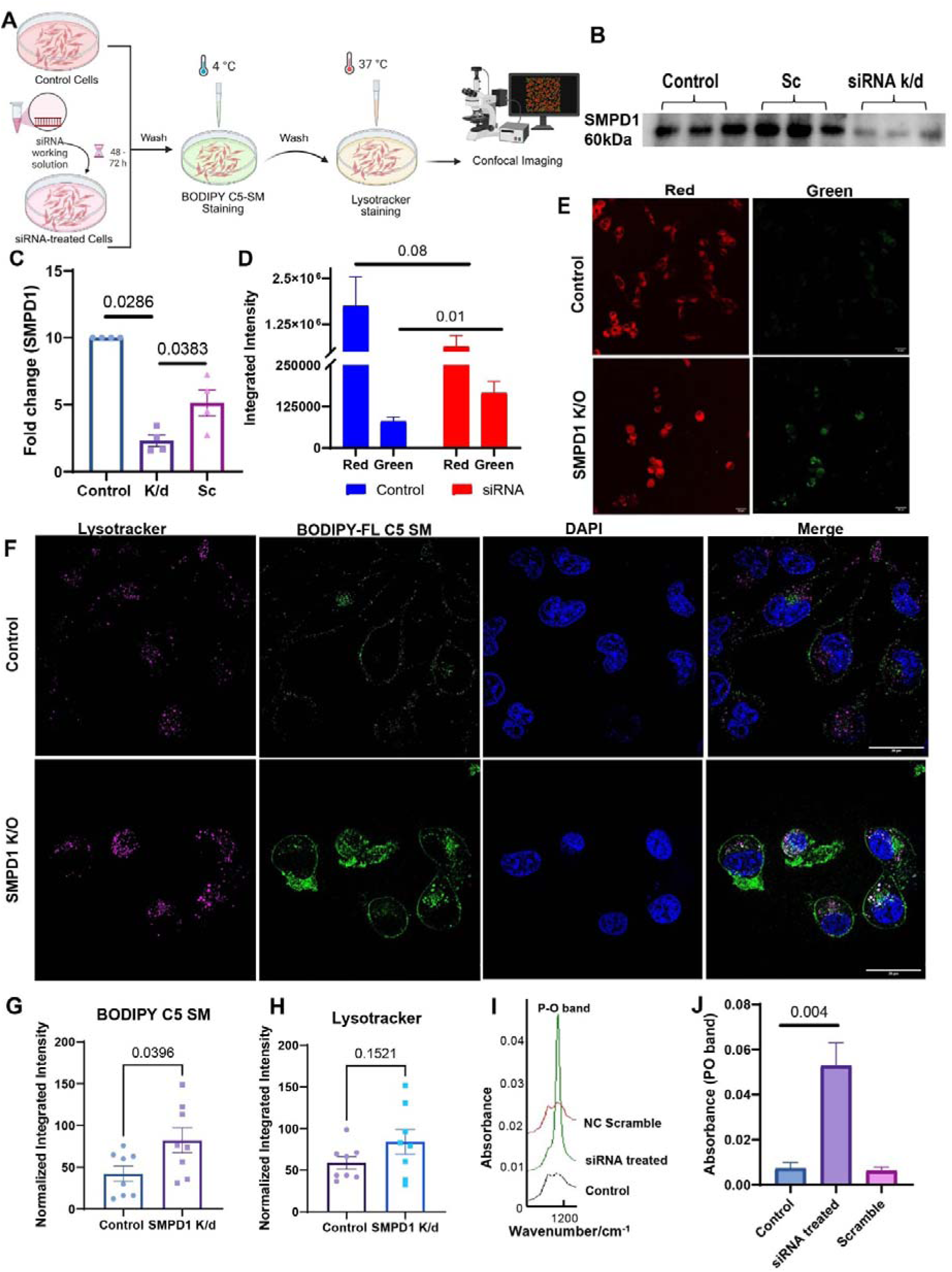
Functional analysis of SMPD1 knockdown on sphingomyelin pathway regulation in SH-SY5Y cells. (A) Schematic representation of the functional analysis of SMPD1 knockdown in SH-SY5Y cells. (B, C) SMPD1 protein levels and mRNA expression were significantly reduced compared with untreated control and Scramble-treated cells. (D) Integrated intensity of the BODIPY C11 561/581 lipid peroxidation dye. (E) Confocal images of control cells and cells treated with SMPD1 siRNA. (F) Immunofluorescence images of control cells and cells treated with SMPD1 siRNA. (G) Integrated intensity of BODIPY C5-SM. (H) LysoTracker signal intensity. (I) FTIR spectroscopy absorbance band corresponding to P-O band. (J) Absorbance ratios. Scale bar= 20µm. (Cntrl= Control, Sc= Scramble, k/d= siRNA knock-down). Data are presented as Mean ± S.E.M, n=4. Statistical significance was determined using Unpaired t-test with p values indicated.

Since a decrease in SMPD1 is expected to alter the ceramide-to-sphingomyelin balance and perpetuate compensatory de novo sphingolipid synthesis, we directly assessed sphingomyelin dynamics using a pulse-chase assay. This approach was chosen because it enables temporal tracking of BODIPY C5 labeled sphingomyelin, allowing separation of changes in clearance rather than relying on steady state measurements. Following pulse labeling and chase, integrated BODIPY C5 fluorescence intensity was significantly higher in SMPD1 knockdown cells (**Figure 5F**), indicating reduced SM catabolism and net SM accumulation. We included LysoTracker staining to assess lysosomal involvement in SM turnover, since SMPD1 encodes an acid sphingomyelinase that functions in the lysosome; altered co-localisation of BODIPY□C5□SM with LysoTracker therefore indicates impaired lysosomal catabolism.

Quantitative analysis confirmed a significant increase in the BODIPY C5 SM signal in the siRNA-treated group (p=0.0396; **Figure 5G**). Furthermore, LysoTracker imaging revealed physically enlarged and more intensely stained lysosomal puncta in the SMPD1 knockdown cells compared to the scramble control (**Figure 5H**). This pronounced morphological shift indicates lysosomal engorgement and dysfunction driven by the failure to efficiently degrade accumulating sphingomyelin. Additionally, we also performed FTIR Spectroscopic analysis of lipid isolated from both control and siRNA-treated group, and observed P-O band was significantly increased in siRNA-treated group (**Figure 5I, Supplementary Figure S7D)**. Since ceramide and Sphingomyelin differ by a phosphocholine headgroup, increased absorbance value for siRNA-treated groups (p=0.004, **Figure 5J)** also strengthens our claim on increase Sphingomyelin turnover after SMPD1 knockdown and is consistent with our immunofluorescence data that reveals a shift toward sphingomyelin accumulation in SMPD1 knockdown group.

### Immunofluorescence analysis validates spatial co-upregulation of SMPD1 and SGMS1 in human PD and murine models

Immunofluorescence analysis was performed on postmortem brain sections from age matched control and PD patients to assess the spatial distribution and intensity of SMPD1 and SGMS1 signals (**Figure 6A**). In human tissue, SMPD1 and SGMS1 immunoreactivity appeared visibly more prominent in PD brain sections compared with AMCs, with stronger signal in neuronal cell bodies, consistent with a global increase in enzyme abundance (inset, **Figure 6B; Supplementary figure S8A)**. Beyond global intensity increases, super-resolution SIM2 analysis revealed a striking subcellular spatial reorganization of both enzymes where a marked, co-localized distribution of both near nuclear rim in separate discrete vacuoles (**Figure 6C**, inset). Quantitative analysis of normalized integrated intensity confirmed this observation, showing elevated SMPD1 signal in PD cases (p=0.0069; **Figure 6D**). Similar profiles were observed in AMCs (inset, **Figure 6E, F; Supplementary figure S8B)**, and quantitative analysis also showed that SGMS1 were also elevated (p=0.03; **Figure 6G**).

**Figure 6:**
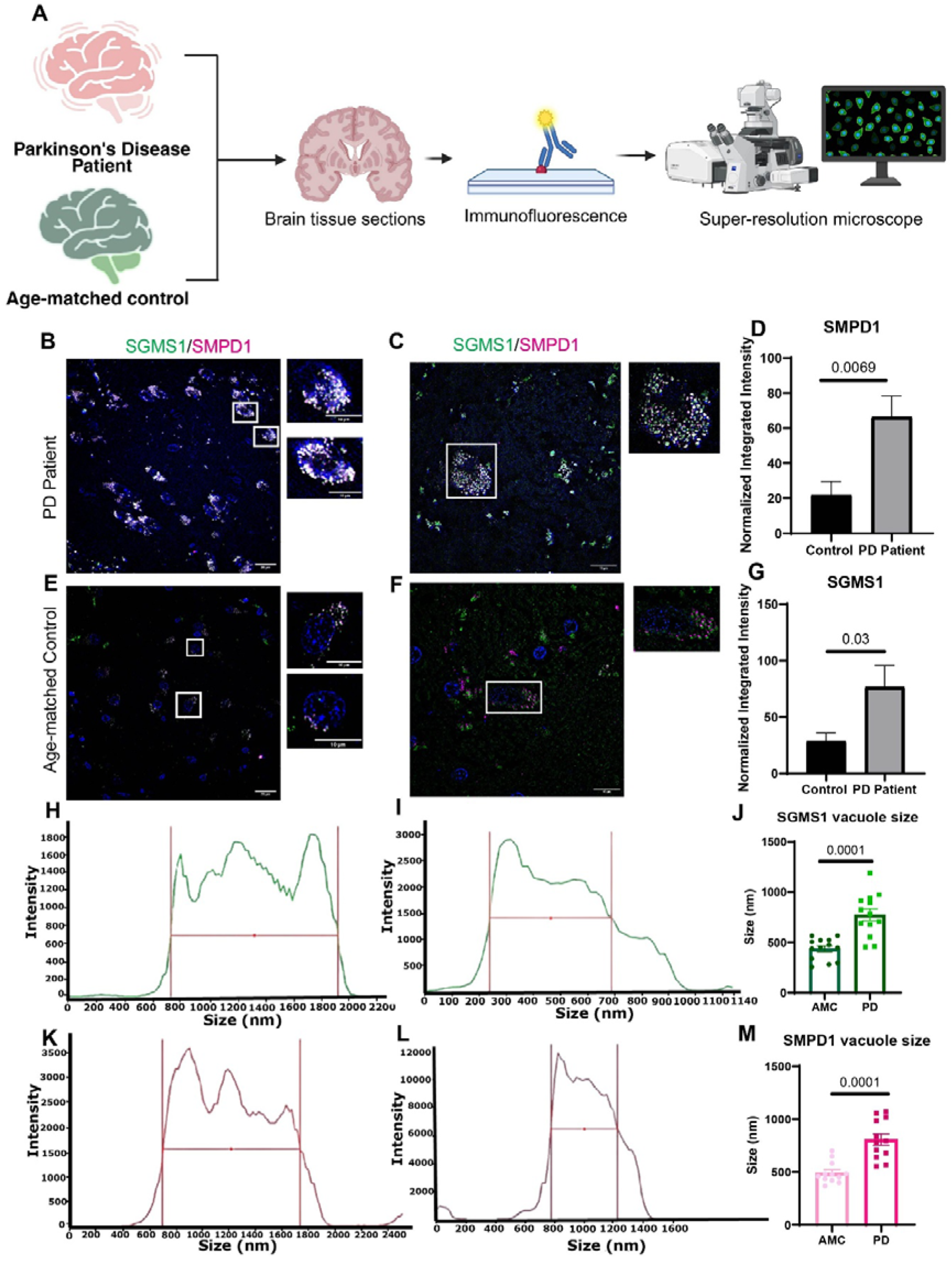
Immunofluorescence analysis of SGMS1 and SMPD1 in human autopsy brain sections. (A) Representative workflow for immunofluorescence images of SGMS1 and SMPD1 in autopsy brain sections from controls and PD patients. (B) Representative immunofluorescence images of SGMS1 and SMPD1 in brain sections from PD patients under 25X and (C) under high 63X magnification. (D) Normalized integrated intensity of SMPD1 in human brain sections. (E) Representative immunofluorescence images of SGMS1 and SMPD1 in brain sections from PD patients under 25X and (F) under high 63X magnification. (G) Normalized integrated intensity of SGMS1 in human brain sections. (H) Line profile diagram showing SGMS1 vacuole in PD and (I) Control. (J) Comparative bar diagram showing difference in SGMS1 vacuole size as observed in PD versus control. (K) Line profile diagram showing SMPD1 vacuole in PD and (L) Control. (M) Comparative bar diagram showing difference in SMPD1 vacuole size as observed in PD versus control. Scale bar= 20µm. Data are presented as Mean ± S.E.M, n=6. Statistical significance was determined using Unpaired t-test, with p values indicated.

High-magnification 3D-SIM2 optical profiling revealed that this observed perinuclear enrichment consists of discrete, spherical vesicular platforms differed in size between PD and AMC groups, for SGMS1 each measuring ∼1000 nm in diameter in PD (**Figure 6H; Supplementary figure S8C**) and ∼450 nm in AMC (**Figure 6I**). Mean SGMS1 vacuole size was 807nm in PD and 450nm in AMC (**Figure 6J**). Similarly, for SMPD1, vacuoles measured ∼1000nm in PD (**Figure 6K**) and ∼450nm in AMC group (**Figure 6L**). Mean SMPD1 vacuole size was 775nm in PD and 437nm in AMC (**Figure 6M**). We observed these microdomains were arranged along the nuclear periphery in a highly ordered, discrete and periodic arrangement (**Supplementary figure S8D)**, with a consistent size difference between the disease and control group, indicating that upregulated SMPD1 and SGMS1 organize into specialized perinuclear lipid-processing platforms during PD pathogenesis. Together, these quantitatively elevated SMPD1 and SGMS1 immunofluorescence level in brain tissue sections is consistent with protein and mRNA findings.

## Discussion

Extracellular vesicles (EVs) are released by most cell types and can be isolated from a wide range of body fluids. EV cargo comprises diverse DNA, RNA, lipid, and protein species that reflect an important part of the parent cell secretome. Although the protein and miRNA content of EVs has been extensively studied, their lipid composition remains less well characterized despite the critical structural and regulatory roles of lipids in EV biogenesis, release, targeting, and uptake. Because EVs can cross the blood-brain barrier, they have emerged as promising minimally invasive sources of diagnostic biomarkers in neurodegenerative diseases, including Parkinson’s disease ^24–28^.

In this study, we successfully isolated and validated plasma-derived small extracellular vesicles (PsEVs) following MISEV guidelines ^21,22^. Crucially, quantitative protein profiling revealed that PsEVs from PD patients were significantly enriched in the canonical tetraspanin CD9, the neuronal adhesion marker L1CAM, and α-synuclein, confirming their utility as a disease-relevant window into central neurodegenerative pathology. After validation of the isolated small extracellular vesicles (sEVs), PsEV-derived lipids were extracted and identified using nano LC-MS/MS.

While dysregulated lipid profiles, including ceramide and sphingomyelin (SM) species, have been previously reported across various tissues in PD, inconsistencies remain, likely due to tissue-specific lipid dynamics. Our approach, focusing on plasma-derived EVs that originate predominantly via endosomal pathways ^29^, makes them an ideal candidate for studying the Ceramide-SM profile and is particularly relevant given the central role of endosomes in ceramide-SM metabolism. Studies have demonstrated a strong correlation between plasma and brain SM levels, underscoring the biological pertinence of analyzing PsEVs lipidomes^9,30,31^.

Ceramides, beyond serving as structural elements for EV membranes, also play critical roles in PD pathophysiology. Mutations in genes encoding sphingomyelin biogenesis proteins are established PD risk factors ^30,31^. We observed a significant 1.6-fold increase in total ceramide levels in PsEVs from PD patients compared to age-matched controls, consistent with previous studies ^32^, accompanied by a reduction in total SM. These findings align with prior reports linking ceramide alterations to disease pathophysiological manifestations ^33^. In line with this, protein levels of sphingomyelin synthase 1 (SGMS1), which catalyzes SM synthesis from ceramide, and sphingomyelin phosphodiesterase (SMPD1, also known as acid sphingomyelinase, ASM), responsible for catabolizing SM back to ceramide ^34^, were elevated in PD patients. Such bidirectional dysregulation of sphingolipid metabolism suggests a dynamic imbalance contributing to altered lipid profiles observed in PD PsEVs.

Crucially, our observation of concurrent SMPD1 and SGMS1 upregulation in both the brain and the gut of rotenone-treated mice aligns with the growing recognition of the gut-brain axis in PD. Because our clinical findings are derived from plasma EVs, which capture a systemic snapshot of cellular health. This dual-organ dysregulation suggests that the ceramide-sphingomyelin metabolic collapse is not restricted to the central nervous system. Rather, it represents a widespread, systemic metabolic stress response that may collectively contribute to the altered circulating vesicle profile

While fragmented evidence from prior studies ^31,32,35^ have noted isolated alterations in ceramide and sphingomyelin (SM) levels in PD, these investigations largely relied on bulk biofluid analyses that lack the critical subcellular spatial resolution required to interpret endosomal-lysosomal lipid dynamics. By leveraging endosome-origin PsEVs, our study advances beyond these observations to delineate a profound, bidirectional disruption of the ceramide-SM axis. Our data clearly establish the systemic ceramide accumulation and concomitant SM depletion are actively driven by the synergistic upregulation of both SGMS1, and the SMPD1. Although earlier reports have detected dysregulated SMPD1 in PD ^36,37^, our orthogonal validation maps its pathological contribution not just as an isolated marker, but as a primary driver of a broader salvage pathway metabolic collapse. This SMPD1-mediated hydrolysis is markedly deleterious, given its role in influencing key PD-relevant cellular processes, including mitophagy, and neuronal survival^35,38^.

By leveraging super-resolution 3D-SIM2 optical profiling, our study provides the first high-resolution visualization of SMPD1 and SGMS1 organizing into discrete, spherical perinuclear vacuoles localized specifically along the nuclear rim. Beyond global enzyme upregulation, we uncovered a striking and novel disease-specific structural feature: while control neurons display small, discrete microdomains measuring ∼437-450 nm, PD neurons exhibit a massive expansion of these perinuclear platforms to ∼775-1000 nm (mean ∼800 nm). This pronounced size doubling and spatial clustering at the nuclear envelope highlight a novel pathological hallmark, demonstrating that hyper-activated sphingolipid metabolism physically restructures nuclear rim microdomains during PD pathogenesis. Dysregulated SMPD also disrupts endosomal trafficking, impairs autophagy, and compromises BBB integrity by affecting endothelial cells, thereby potentially exacerbating neurodegenerative mechanisms ^39^. Furthermore, the concurrent upregulation of the SM lysosomal transporter ABCA5^32^ corroborates this mechanism, perfectly aligning with our observed SM depletion and ceramide accumulation.

Recent work by Peng et al.^40^ also provides an important translational support for the biomarker potential of sphingolipid abnormalities in Parkinson’s disease, particularly through the analysis of plasma ceramides and glycosphingolipid ratios. However, the absence of mechanistic implications of the findings require validation in longitudinal cohorts and functional studies. In contrast, our study delineates the sphingolipid pathway more comprehensively, integrating sEV lipidomics, SMPD1-based functional validation, and tissue-level confirmation to support a model of impaired lysosomal sphingolipid degradation accompanied by compensatory ceramide synthesis. Collectively, our findings extend beyond isolated lipid correlations to reveal a spatially resolved collapse of SMPD1-dependent sphingolipid homeostasis in PD pathophysiology.

Additionally, we observed the sphingolipid salvage pathway was dysregulated as part of a multi-pathway metabolic collapse. SPTLC mRNA levels remained unchanged, ruling out reduced de novo synthesis as the primary defect. Instead, SM downregulation is driven by SMPD1 upregulation (enhanced hydrolysis), while ceramide accumulation (C16:0, C18:0, C18:1, C20:1) results from SM hydrolysis by SMPD1, and metabolic collapse of salvage pathway. Ceramide synthase (CerS5) upregulation represents a compensatory response to sphingosine accumulation caused by ASAH1 upregulation and SphK1 downregulation, elevating ceramide level. The critical bottleneck is SphK1 downregulation, which blocks sphingosine to S1P conversion^41,42^, depleting pro-survival signaling and causing ceramide/sphingosine rheostat imbalance toward dysregulation^43,44^. SGPL1 downregulation further impairs autophagy via PE depletion, directly linking sphingolipid metabolism to α-synuclein accumulation ^45^. Our pulse-chase experiments showed clear accumulation of BODIPY-SM in the siRNA-treated group, with stronger relative fluorescence compared with the evenly distributed signal observed in controls indicating accumulation of labeled sphingomyelin. The concomitant increase in LysoTracker signal in the same group suggests that SM accumulation following SMPD1 knockdown may provoke a compensatory lysosomal response. In addition, BODIPY 581/591 C11 lipid peroxidation analysis revealed a marked red-to-green fluorescence shift in siRNA-treated cells, indicating elevated oxidative membrane stress that was concordant with the lipidomic signature of ceramide enrichment and sphingomyelin depletion. Consistent with biochemical data, FTIR spectroscopy of lipid extracts from SMPD1 knockdown cells showed increased P-O band absorbance, providing additional support for enhanced sphingomyelin turnover in SMPD1 knock-down condition.

Interestingly, we observed that both clinical SMPD1 hyperactivity and SMPD1 knockdown lead to a collapse of the Ceramide: Sphingomyelin ratio. In the clinical state, chronic upregulation of SMPD1 led to an acute accumulation of ceramides whereas in experimental SMPD1 knockdown, the failure to degrade sphingomyelin led to lysosomal engorgement and a metabolic bottleneck. These two extremes represent a ‘U-shaped’ homeostatic curve where any deviation, whether through overactivation or silencing arrests the salvage pathway, resulting in PINK1 downregulation and the subsequent arrest of mitophagy. Collectively, both processes demonstrate that disrupting the sphingomyelin-ceramide balance whether through clinical overactivation or modeled genetic knockdown leads to an identical convergence of lysosomal storage pathology, systemic lipid peroxidation, and mitophagy arrest, establishing sphingolipid homeostatic collapse as a central driver of Parkinson’s disease pathogenesis.

Crucially, our cross-disease comparison reveals that this is not a generic feature of neurodegeneration. While Alzheimer’s disease cohorts also exhibited elevated SMPD1, this was accompanied by a downregulation of SGMS1, representing a unidirectional collapse of sphingomyelin synthesis. In contrast, PD PsEVs demonstrate a unique, bidirectional disruption. The concurrent upregulation of SGMS1 in PD likely represents a failed compensatory effort by the cell to manage the ceramide accumulation driven by SMPD1 hyperactivity. Because SMPD1 upregulation outpaces SGMS1, this compensatory loop is overwhelmed, forcing the ceramide burden into the salvage pathway and precipitating the downstream autophagic failure. Together, SMPD1-driven hydrolysis, causes ceramide accumulation which not only converges to mitophagy failure^23^ but also paves way for salvage pathway collapse, which ultimately depletes pro-survival S1P signaling thereby exacerbating lipid homeostatic collapse.

Importantly, our study highlights distinct alterations in sphingolipid metabolism associated with the postural instability and gait difficulty (PIGD) clinical subtype of PD. Expression levels of both SGMS1 and SMPD1 enzymes were significantly reduced in PsEVs from patients with PIGD compared to the tremor-dominant (TD) subtype. This differential enzyme expression suggests subtype-specific lipid metabolic remodeling, which may underpin the more aggressive clinical progression and poorer prognosis characteristic of PIGD ^46,47^. The distinctive lipid enzyme signature in PIGD underscores the heterogeneity of PD pathophysiology ^48^, and supports the utility of PsEVs lipid profiling to capture clinically meaningful molecular differences.

The study has some limitations. First, it is cross-sectional study, with all patients recruited from a single tertiary neurology outpatient clinic, which may limit the external validity of the findings. Second, although blood samples were obtained under fasting conditions, most patients were already receiving levodopa and other dopaminergic therapies at the time of collection, which may have influenced circulating EV and lipid profiles. In addition, the sample size is relatively modest, particularly for subgroup analyses, and stratified results should therefore be interpreted with appropriate caution. Beyond elucidating the molecular interplay of SMPD1 dysregulation as a driver of Parkinson’s disease pathogenesis, large multicentre, population-based longitudinal cohorts, with explicit stratification of drug-naive and medicated patients, will be pivotal to establish the specificity and temporal behaviour of plasma sEV-associated sphingolipid signatures.

Taken together, our data reveals holistic representation of the sphingomyelin metabolic network, capturing changes across de novo synthesis, sphingomyelin synthesis, hydrolysis, and ceramide salvage pathways rather than focusing on a single enzymatic node. By combining plasma EV lipidomics, enzyme expression at the protein and mRNA levels, FTIR based chemical signature, and high-resolution SIM2 immunofluorescent profiling in human autopsy provides a discrete spatial organization of SMPD1/SGMS1 in immunolabelled autopsy brain sections. Orthogonal validation in in vitro, in vivo, transgenic, and human clinical samples further supports the translational relevance of these findings and suggests that SMPD1-centered sphingolipid dysregulation is a critical feature of PD pathobiology with potential biomarker and therapeutic implications. Finally, the differences observed across clinical PD subtypes, together with the differential expression of these enzymes in other neurodegenerative diseases such as AD and MS, underscore the importance of considering clinical subtype when interpreting lipid alterations. These findings also suggest that the observed changes are disease-specific rather than a generic feature of neurodegeneration, with potential implications for mechanistic understanding and the development of personalized therapeutic strategies.

## Methods

### Subject recruitment

Peripheral blood samples were collected by venipuncture from patients with Parkinson’s disease (n = 90) and age-matched controls (n= 60) at the All India Institute of Medical Sciences, New Delhi (Table 1). Parkinson’s disease was diagnosed according to the Movement Disorder Society clinical diagnostic criteria for Parkinson’s disease ^49,50^. Patients were further classified as tremor-dominant (TD), postural instability/gait difficulty (PIGD), or indeterminate subtypes using the Movement Disorder Society-Unified Parkinson’s Disease Rating Scale (MDS-UPDRS), following the method described by ^51^. Briefly, tremor and PIGD scores were calculated from predefined items, and a tremor-to-PIGD ratio was derived; TD was defined as a ratio of ≥1.5, PIGD as ≤1.0, and indeterminate as 1.0-1.5. All participants provided written informed consent, and the study was approved by the Institutional Ethics Committee of the All India Institute of Medical Sciences, New Delhi, India (Ref. No.: IECPG-766/30.11.2022).

**Table 1.** Demographic Details of the Subject Recruited.

| Parameters | Age-matched Control | Parkinson's disease patient | P Value |
| --- | --- | --- | --- |
| Number | 60 | 90 | - |
| Age (Mean ±SD) | 52.5 ±9.364 | 54.426 ±10.415 | 0.2403 |
| Male | 48 | 61 | 0.1447 |
| Female | 12 | 29 | 0.1447 |
| NTA (Mean ±SD) | 1.51E+11 ±2.14 | 4.49E+11 ±4.79 | 0.0001 |
| α-synuclein (pg/mL) | 181.9 ±93.99 | 886.6 ±388.1 | 0.0001 |
| Age at Onset (Years) | - | 53.64 ±11.19 | - |
| Disease Duration (Years) | - | 4.42 ±4.18 | - |
| HY Score (Range) | - | 1 - 4 | - |
| TD | - | 48.9% | - |
| PIGD | - | 44.4% | - |

### Sample processing

Plasma was separated from whole blood by centrifugation at 2,500 × g for 10 min at 4°C, followed by an additional spin at 10,000 × g for 30 min to remove microvesicles. The resulting clarified plasma was aliquoted and stored at −80°C. Prior to analysis, samples were thawed and centrifuged again at 3,000 × g for 5 min at 4°C. Peripheral blood mononuclear cells (PBMCs) were isolated by density-gradient centrifugation using Histopaque 1077 (Sigma), followed by red blood cell lysis with RBC lysis buffer (11814389001, Roche).

### Isolation of sEVs

Plasma samples (500 μL) were diluted with 500 μL phosphate-buffered saline (PBS, pH 7.4), subjected to ultrafiltration, and transferred to Ultra-Clear ultracentrifuge tubes (Beckman Coulter). The samples were then centrifuged at 120,000 x g for 90 min at 4 °C using a TLA-120 fixed-angle rotor (Beckman Coulter) to pellet extracellular vesicles. The supernatant was carefully removed, and the pellet was resuspended in 200 μL PBS. A second ultracentrifugation step at 120,000 x g for 90 min at 4 °C was performed to further enrich the EV fraction. The final pellet was resuspended in 100 μL PBS and used for nanoparticle tracking analysis (NTA). In addition, small extracellular vesicles (sEVs) were isolated from human plasma samples using chemical precipitation combined with ultrafiltration, as described previously^24,52^.

### Transmission electron microscopy

Transmission electron microscopy (TEM) was used to examine the ultrastructural morphology of plasma-derived small extracellular vesicles (PsEVs). PsEVs were diluted 1:100 in 1× PBS and adsorbed to carbon-coated copper grids (Ted Pella, 01843) for 30 min at room temperature. The grids were then blotted dry, negatively stained with 1% aqueous uranyl acetate for 15 s, blotted again, and examined using a transmission electron microscope (Talos S, Thermo Scientific, USA).

### Nanoparticle tracking analysis

Nanoparticle tracking analysis (NTA) was used to assess particle size distribution and concentration in solution. In this study, measurements were carried out using the ZetaView system (Particle Metrix, Germany) in scatter mode. PsEVs were diluted 1:1000 in 1× PBS, and 0.5 mL of the diluted sample was introduced into the NTA chamber for analysis. Three measurement cycles were acquired by scanning 11 positions per cycle, with 60 frames recorded at each position under high video settings, using autofocus, camera sensitivity of 80.0, shutter speed of 150, scattering intensity of 5.0, an integrated 488 nm laser, and a chamber temperature of 25°C. Videos were captured using a CMOS camera and analyzed with ZetaView software version 8.05.12 using preset parameters of maximum particle size 1000, minimum particle size 10, and minimum particle brightness 30.

### Western blot

PsEV samples were normalized to the initial biofluid input volume before western blot analysis. The sEV-enriched fractions were mixed with sample loading dye, and equal volumes of 20 μL were loaded onto a 12% SDS-PAGE gel. After electrophoresis, proteins were transferred in wet mode using a Bio-Rad western blot system onto a 0.22 µm PVDF membrane. The membrane was then blocked with 3% BSA in Tris-buffered saline containing 0.1% Tween 20 (TBST). Primary antibodies were diluted 1:5000 in 1.5% BSA/TBST and incubated overnight at 4°C, including anti-CD9 (PA5-86534, Invitrogen), anti-Flotillin-1 (PA5-17127, Invitrogen), anti-Parkin (PA5-13399, Invitrogen), anti-SMPD1 (A23894, AbClonal), anti-SGMS1 (A15008, AbClonal), anti-TH (A5079, AbClonal), and anti-β-actin (AC038, AbClonal). The blot was developed using HRP-based chemiluminescence with the Femto LUCENT™ PLUS-HRP kit (G-Biosciences).

### RNA isolation and gene expression assays

Total RNA was isolated from tissues and cells using the RNeasy Mini Kit (74124, Qiagen) following the manufacturer’s protocol. Complementary DNA was synthesized using the RevertAid First Strand cDNA Synthesis Kit (K1622, Thermo) following the manufacturer’s protocol. Quantitative real-time PCR (qRT-PCR) was performed using PowerUp™ SYBR™ Green Master Mix (A25742, Applied Biosystems) and Pre-designed TaqMan® Gene Expression Assays (4331182, Thermo) on a qTOWER³ Real-Time PCR System (Analytik Jena). Relative gene expression was quantified using the comparative Ct method (ΔΔCt Method).

### Lipid isolation

For lipid isolation modified Folch method was used ^53^, which employed a biphasic solvent system composed of chloroform and methanol in a 2:1(v/v) ratio. PsEVs samples were thawed on ice prior to addition of 200μl of ice-cold chloroform (Sigma) to 20μl PsEVs and then vortexed for 20 seconds. Next, 200μl of ice-cold Methanol (Sigma): 0.9% saline (NaCl) solution in a 1:1(v/v) ratio was added to induce phase separation and then vortexed rigorously for 20 seconds. Finally, tubes were centrifuged at 18,000 × g for 20 minutes and organic bottom layer was collected, and transferred into a fresh polypropylene tube and analysed by LC-ESI MS/MS.

### LC-MS conditions

The LC-MS analysis was performed on a high resolution OrbitapExploris 120 mass spectrometer (Thermo Fisher Scientific) equipped with a Vanquish UHPLC (Thermo Fisher Scientific). An aliquot of 10µL of the sample was injected into the LC/MS/MS instrument using an auto-sampler. The Chromatographic separation was carried out using the Hypersil Gold C18 column (100mm length*2.1mm internal diameter, 1.9 Micron Particle size: Thermo Scientific.), column temperature kept as 50, auto-sampler temperature kept as 10. A two-component mobile phase consisting 0.1% formic acid in Millipore water (Component A) and 100% methanol (Component B) at 0.3mL/min flow rate was used. The gradient program was at 0 min, 5 %B; at 2 min, 5 %B; at 15 min, 95 %B; at 17 min, 95 %B; at 18 min, 5 %B; at 22 min, 5 %B; The total run time was 22 min.The Heated-electrospray ionization (H-ESI) under positive and negative mode (polarity switching) was used as anionisation source of mass spectrometry. The typical MS conditions were: capillary voltage was set at 3500V, 2500V for positive and negative ionisation modes respectively, nitrogen was used as sheath gas and auxiliary gas, and their flow rates was maintained at 60 (arbitrary value) and 12 (arbitrary value), respectively. The ion transfer tube and vaporizer temperatures were maintained at 300 and 330 respectively; the CID gas pressure was set at 1.5 mTorr. The Q1 resolution was 0.7 (FWHM). The mass spectrometer was scanned in Full scan and as well as Data Dependent MS2 (ddMS2). The full scan parameters are scanning range (*m/z*) 100-1500, Orbitap resolution 60,000 (FWHM), RF lens (%) 70, Data type Profile. The ddMS2 parameters are isolation window (*m/z*) 1, collision energy type normalized, HCD collision energies (%) 20, 30, 60. Orbitap resolution 15,000 (FWHM), Data type Centroid, peak filters; intensity threshold 1.0e5, charge state singly charged, dynamic exclusion i) PPM= ±5, ii) exclusion duration = 5 sec.

### Identification of metabolites and data processing

Untargeted metabolite and lipid screening of the acquired data was performed using Compound Discoverer software (v. 3.3 SP3; Thermo Fisher Scientific). Metabolite identification relied on retention time alignment, accurate mass, and MS/MS spectral matching against ChemSpider, mzCloud, mzVault, MassList, KEGG, and HMDB databases. The resulting data matrix was stringently filtered by applying peak intensity cut-offs, evaluating chromatogram peak quality, and establishing fold-change thresholds to yield the final identified and quantified metabolite and lipid profiles.

### Annotation of lipid classes and species

Glycerophospholipids are referred to as phosphatidic acids (PA), phosphatidylinositols (PI), phosphatidylserines (PS), phosphatidylglycerols (PG), phosphatidylethanolamines (PE), phosphatidylcholines (PC); Polyketides (LMPK); Glycerolipids as Diradylglycerolipids (DG) and Triradylglycerolipids (TG); and Sphingolipids as Ceramides (Cer) and sphingomyelins (SM) ^54,55^. Lipid species and subspecies are annotated based on their molecular composition using the following format indicated in (Table 2).

**Table 2.**
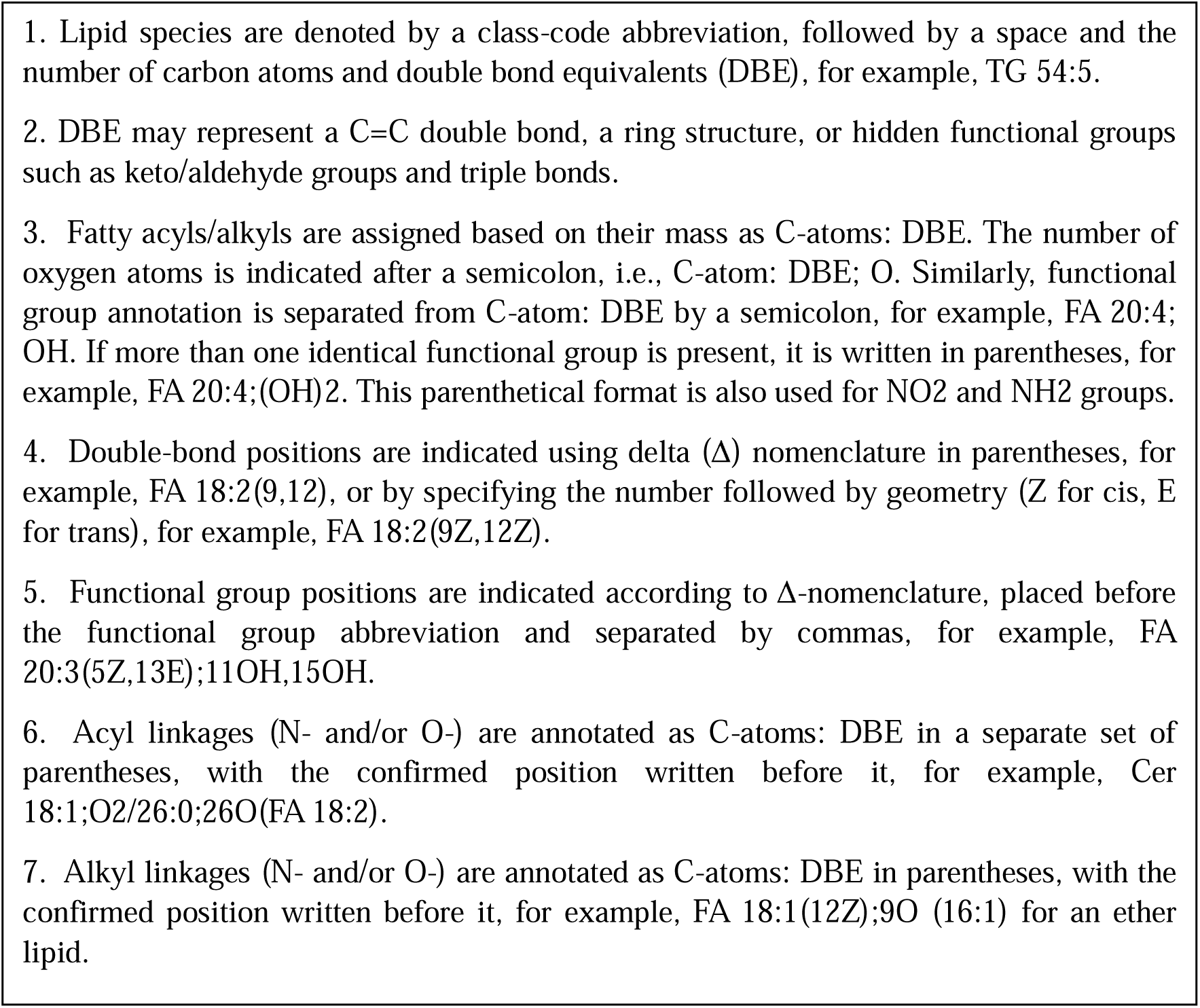
Nomenclature of Lipid Species.

**Table 3.**
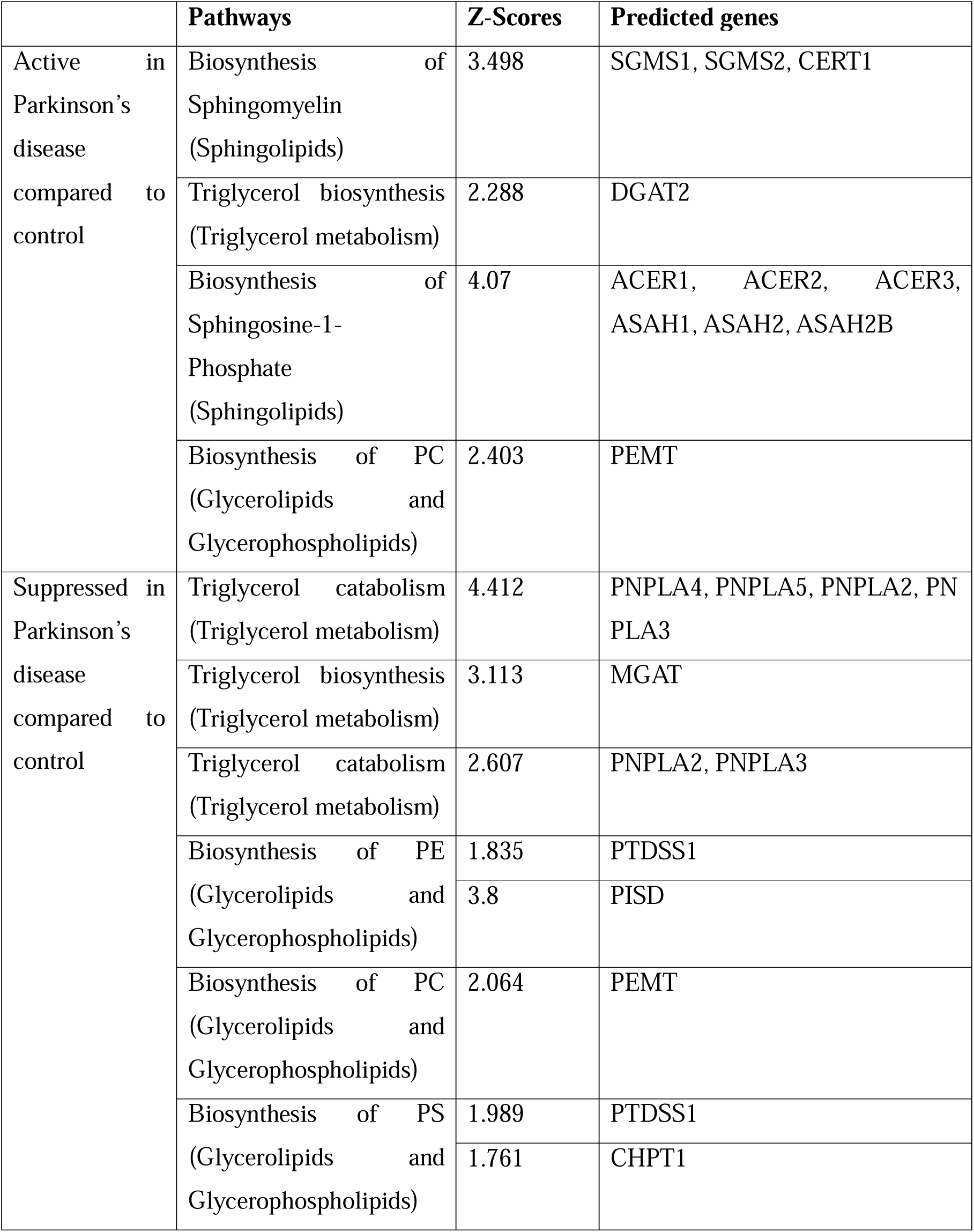
summarizes all the pathways activated or suppressed in Parkinson’s disease.

### Immunofluorescence staining of Brain tissue

Human autopsy brain samples were obtained from the NIMHANS Brain Bank. Brain sections underwent deparaffinized in xylene and rehydrated through graded ethanol washes of 100%, 90%, 75%, 50%, and 30%, followed by a final rinse in 1x PBS, with each step duration of 5 minutes. The sections were incubated with primary antibodies against CD9 (Catalog # PA5-86534, Invitrogen), SMPD1 (ab181372, Abcam and OTI3H7, Invitrogen), and SGMS1 (PA5100331, Invitrogen), followed by TRITC-conjugated secondary antibodies (Goat anti-Rabbit IgG (H+L), Catalog # A11008 and A11032). After 120 min at room temperature, the sections were washed three times with 0.1% PBST, 5 min per wash, mounted, and visualized using a Zeiss Elyra SIM² microscope.

### Pulse-chase experiment to study sphingolipid metabolism

1×10^4^ SH□SY5Y cells were seeded in chambered glass-bottom plates (C4SB-1.5H, Cell vis). Following day, adherent cells were washed with HBSS/HEPES buffer at room temperature to remove residual growth medium and non□viable cells. 300 µL of BODIPY FL C5□sphingomyelin working solution (1 mM stock) prepared as described in ^56^ was then added to fully cover the cell growth area, and were incubated at 4□°C for 30□min in the dark. For negative controls, HBSS/HEPES buffer was used in place of the BODIPY FL C5□sphingolipid working solution during this incubation. After the 30□min incubation at 4□°C, the working solution was completely removed, and cells were washed three times with ice□cold DMEM/F12 medium. Fresh media at room temperature was then added to the cells, and cells were counterstained with Hoechst (33342, Thermo) for nuclear staining and LysoTracker (L12492, Thermo) for lysosomal staining which was followed by incubation at 37□°C with 95% humidity and 5% CO□ for 30□min, in dark. Cells were subsequently washed with fresh media at room temperature. Stained cells were imaged directly using a Zeiss Elyra SIM² microscope at 63× magnification.

### FTIR Analyses

FTIR spectra were recorded using an Agilent Cary 600 spectrometer fitted with a DTGS detector and an ATR accessory. 10µL of each sample were applied to the diamond ATR crystal and analyzed at room temperature, with chloroform used as the background. To evaluate reproducibility and sample heterogeneity, three independent FTIR measurements were acquired for each lipid sample. All spectra were visually examined, showing minor within-group variation but no outliers. In total, 9 spectra (3 samples × 3 measurements) from the untreated control, siRNA-treated, and scramble-treated groups were included in the analysis. ATR crystal was cleaned after each run with ethanol and lint-free wipes, and the instrument was continuously purged with dry air, as per established protocols^57,58^. Spectral data were acquired in the range of 800-4000 cm^-^¹, with 128 scans averaged for each measurement. Data analysis was carried out using Agilent ResolutionPro, GraphPad Prism, and the Kinetics toolbox in MATLAB. Second-derivative spectra were generated after applying 4 cm^-^¹ smoothing using Erik Goormaghtigh’s Kinetics software, and the spectra were carefully examined to eliminate water-vapor artifacts. ^59^. The amide I region (1700-1600 cm^-^¹) was curve-fitted following baseline subtraction, with initial peak positions determined from the second-derivative spectra. Final fitting was performed using Voigt functions, with the baseline iteratively optimized by least-squares regression. Peak deconvolution employed Voigt line□shape functions to capture both Gaussian and Lorentzian broadening, yielding lower residuals and better representing asymmetric bands in the amide and lipid regions. Baseline correction used a six□point procedure with reference wavenumbers at 3200, 2700, 1950, 1485, 1000, and 801□cm□¹ to standardize baseline fluctuations and ensure consistent representation of chemical information.

### Statistical analysis

To ensure robust analysis, only lipid species detected in all samples were retained for statistical and bioinformatics evaluation using the free web-based software MetaboAnalyst (version 5.0). Samples were median-normalized, log10-transformed, and auto-scaled. Lipids were considered statistically significant candidate features when they met a fold-change threshold of −2 ≤ Fold Change ≤2 and a p-value of ≤0.05. In MetaboAnalyst 5.0, Student’s t-test, volcano plots, ANOVA, PCA, heatmaps, and box-and-whisker plots were generated. Statistical analyses were also performed in GraphPad Prism 8.0 using the Unpaired Student’s t-test, with significance set at p < 0.05.

## Acknowledgement

The authors acknowledge Indian Council of Medical Research (ICMR) for providing financial support. All graphical illustrations were created using BioRender and images were prepared in Adobe Illustrator. We thank Dr. Sarika Gupta (National Institute of Immunology) for providing Parkin□knockout mice. We are grateful to the Human Brain Bank, NIMHANS, for providing autopsy brain tissue samples from clinically diagnosed PD patients and age matched controls for immunofluorescence staining. We also acknowledge SAIF AIIMS SRM Facility for access to the Zeiss Elyra SIM² microscope.

## Consent Statement

All human subjects provided informed consent.

## Data Availability Statement

Supplementary datasets are available from the corresponding author upon reasonable request. All data necessary to evaluate the conclusions of this paper are provided in the Supplementary file, which is available with this article

