## Supplementary Information for "Plasma Extracellular Vesicle Lipidomics Reveals an SMPD1-Driven Sphingomyelin Salvage Pathway Collapse in Parkinson’s Disease"

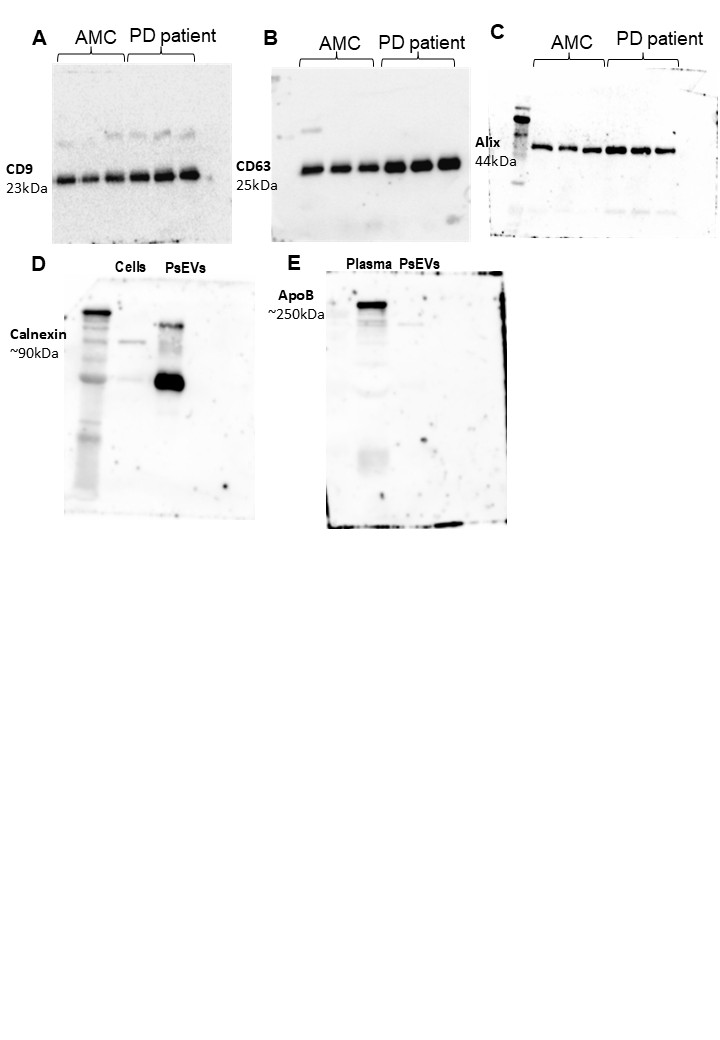

**Figure S1:** Uncropped Western blot of EV markers: CD9 (A), CD63 (B), Alix (C), Calnexin (D) and Apolipoprotein (E).

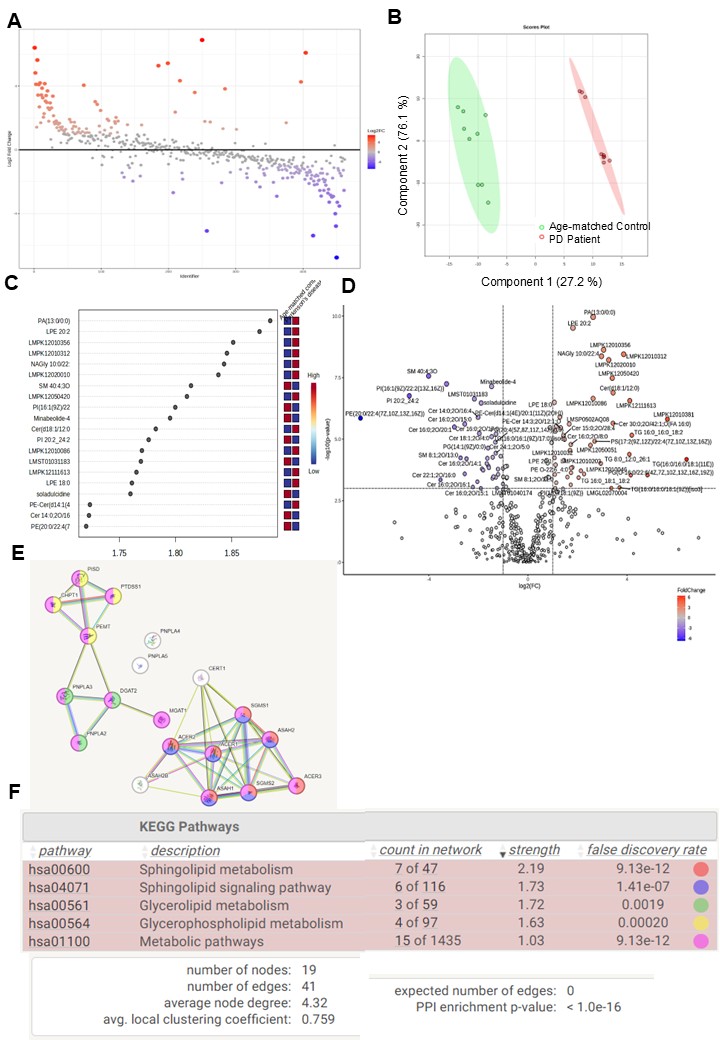

**Figure S2:** Graphical illustration of the total differentially regulated lipid species based on their fold Change (A). Volcano plot for differentially expressed lipid species (B). PLS-DA plot (C) plotted based on Variable importance plot (VIP) showing top 20 differentially expressed lipids (D), String analysis of the predicted genes (E). KEGG pathway and functional enrichment of the predicted genes (F).

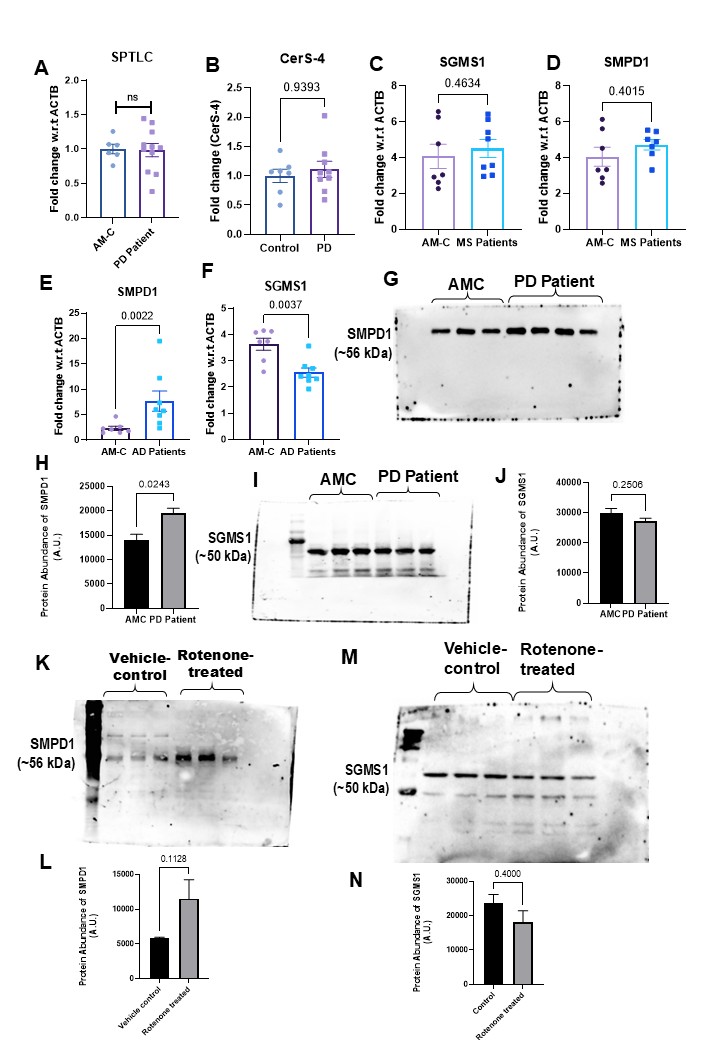

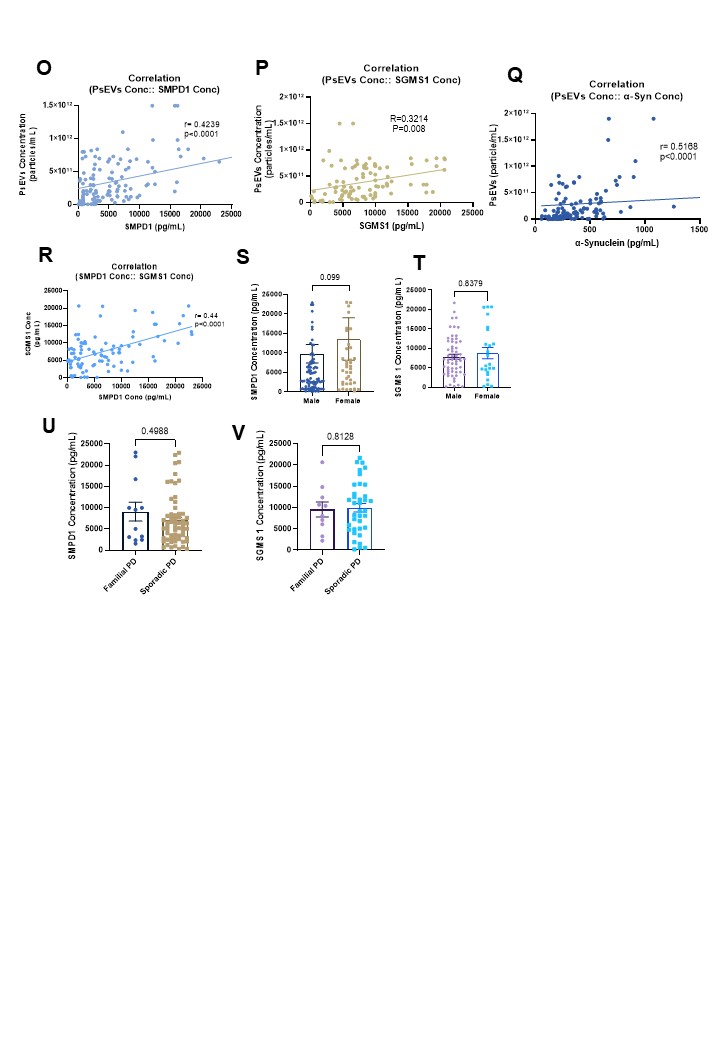
**Figure S3:** Total mRNA expression levels observed between PD patients and controls for SPTLC (A), CerS-4 (B), in AD patients SGMS1 (C), SMPD1 (D). Similarly in MS patients SMPD1 (E), SGMS1 (F). Uncropped Western blot of SMPD1 (G-H) and SGMS1 (I-J) in Clinical samples. Similarly for SMPD1 (K-L) and SGMS1 (M-N) in mice brain lysates. Correlation between PsEVs Concentration- and SMPD1 Concentration (O), SGMS1 (P), α-synuclein concentration (Q) and Correlation between SMPD1 and SGMS1 concentration (R). Gender-wise differences in concentration of SMPD1 (S) and SGMS1 (T). Difference between Familial and Sporadic PD cases in concentration of SMPD1 (U) and SGMS1 (V).

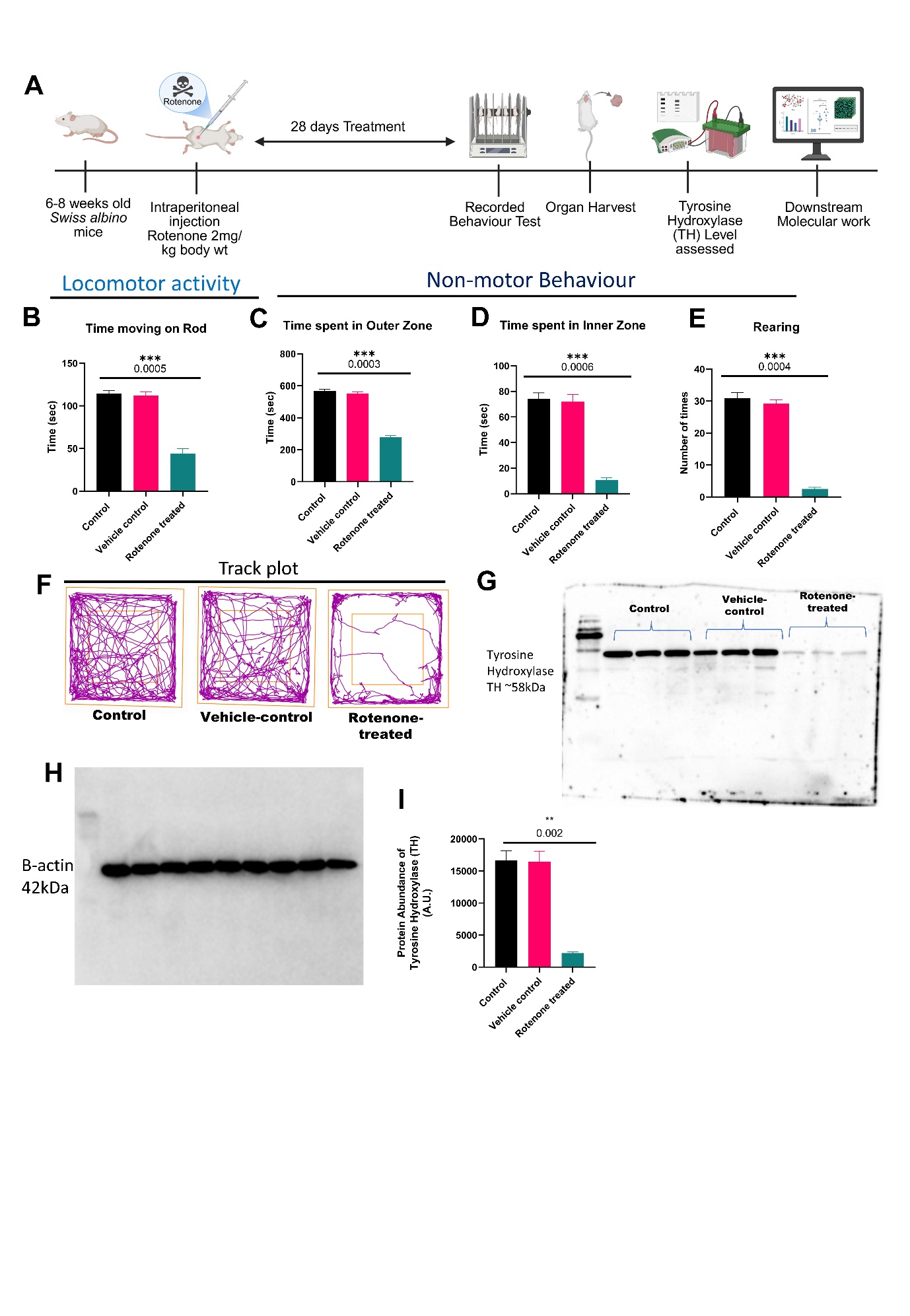

**Figure S4:** Development of PD model in Swiss albino mice (A), Locomotor activity measured by Rotarod (B), Non-motor behaviour (anxiety) measured by Open Field Test (OFT)- Time spent in outer zone (C), Time spent in Inner Zone (D), No. of Rearing times (E) and Track plot (F), Western blot of TH (G) and β-actin (H), Densitometry analysis for TH (I).

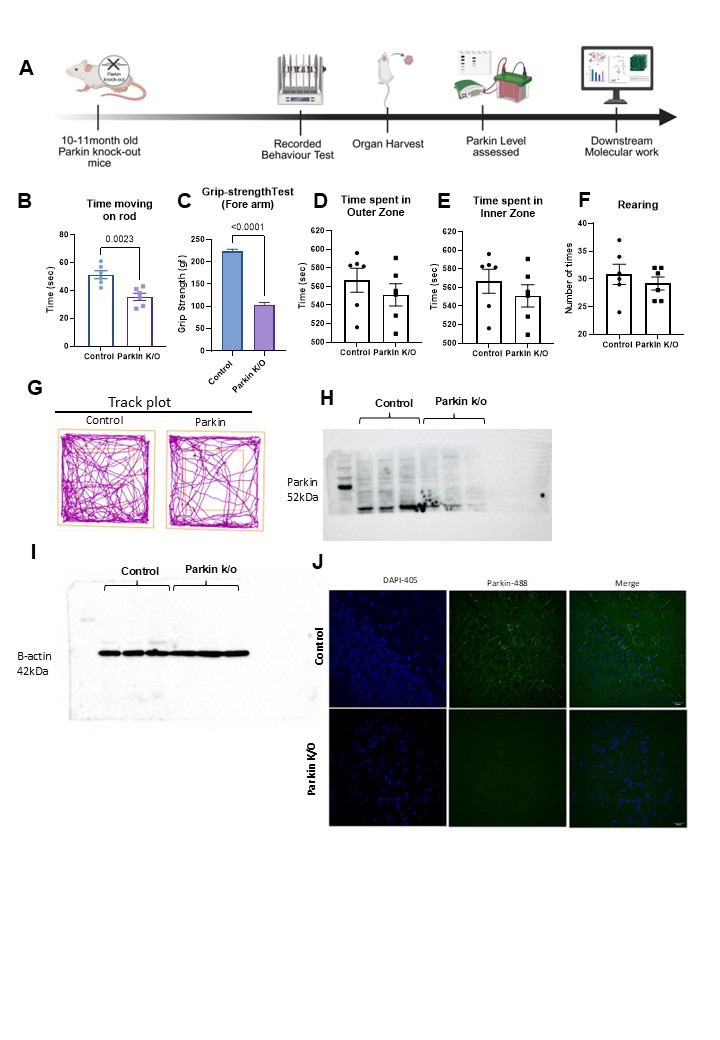

**Figure S5:** Graphical Representation of workflow (A), Locomotor activity measured by Rotarod (B), Grip-strength test (C), Non-motor behaviour (anxiety) measured by Open Field Test (OFT)- Time spent in outer zone (D), Time spent in Inner Zone (E), No. of Rearing times (F) and Track plot (G), Western blot of Parkin (H), and Beta-actin (I), Immunofluorescence image of Parkin in brain-section (J) Scale bar (20 µm).

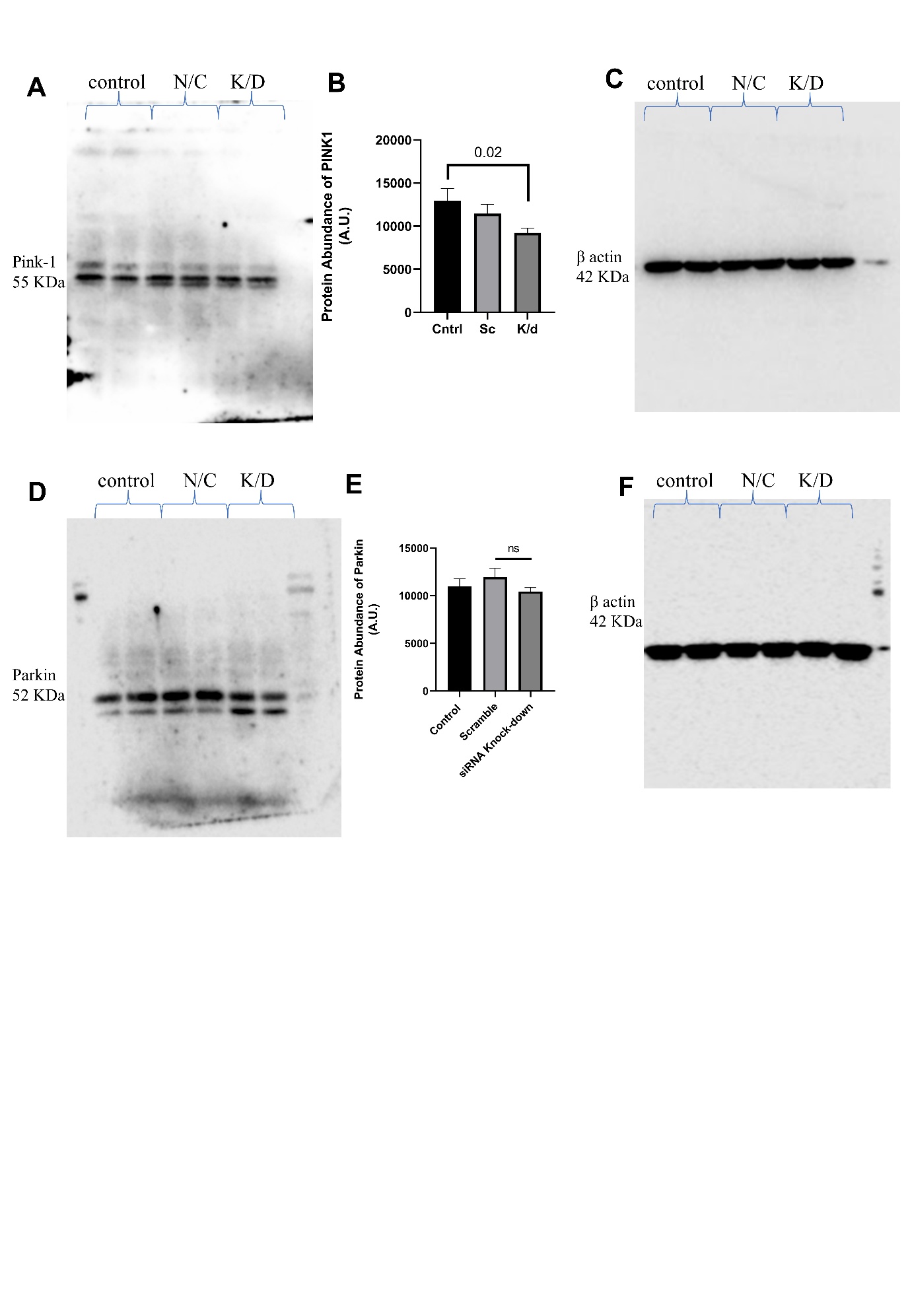

**Figure S6:** Uncropped Western blot of PINK1 (A), Normalised Densitometry analysis of PINK1 (B), Uncropped Loading control β-actin (C), Uncropped Western blot of Parkin (D), Normalised Densitometry analysis of Parkin (E), Uncropped Loading control β-actin (F).

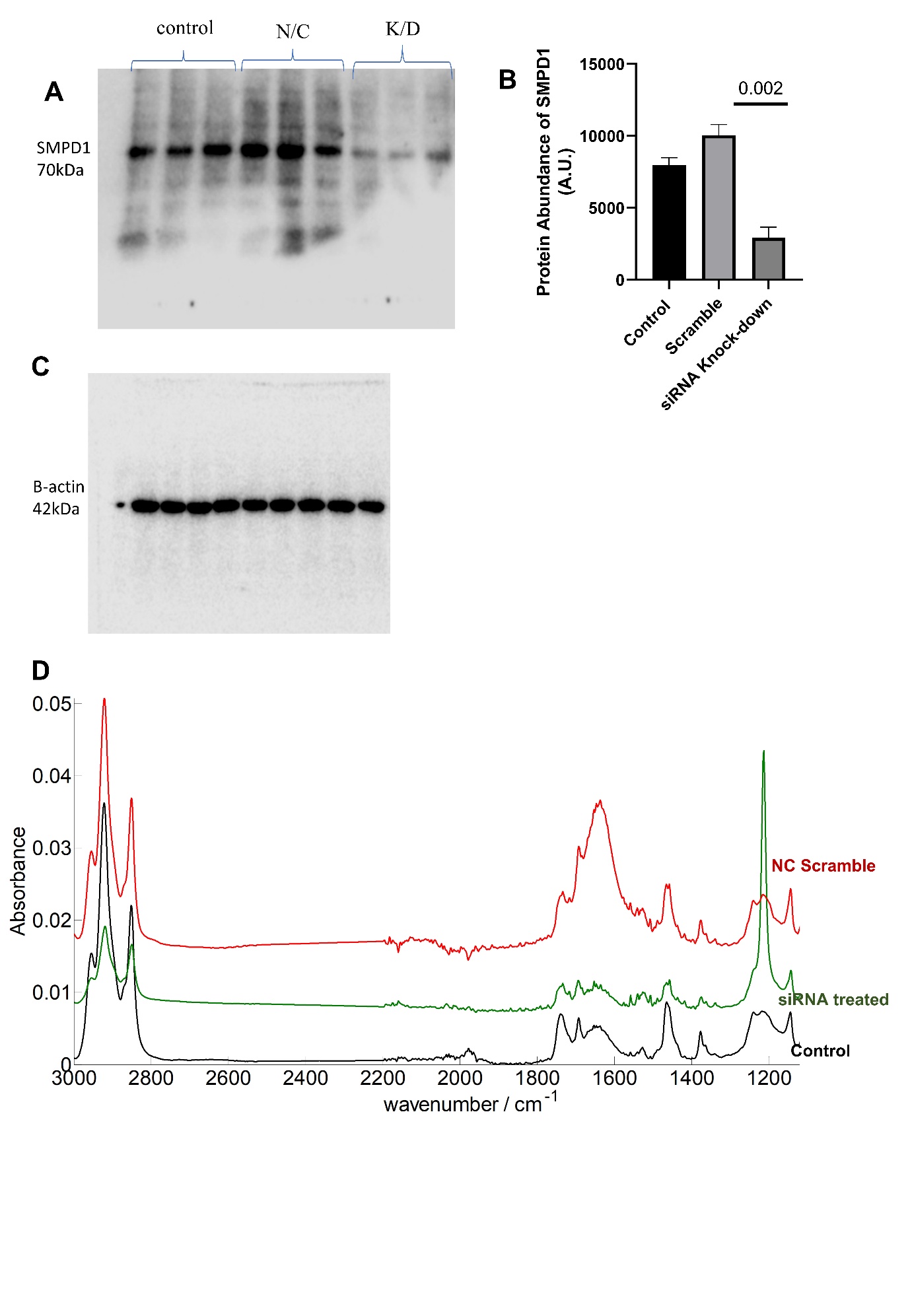

**Figure S7:** Uncropped western Blot image for SMPD1 (A), and densitometry analysis (B). Uncropped western Blot image for β-actin (C). FTIR Spectra of isolated lipid from control and siRNA- treated cells (D).

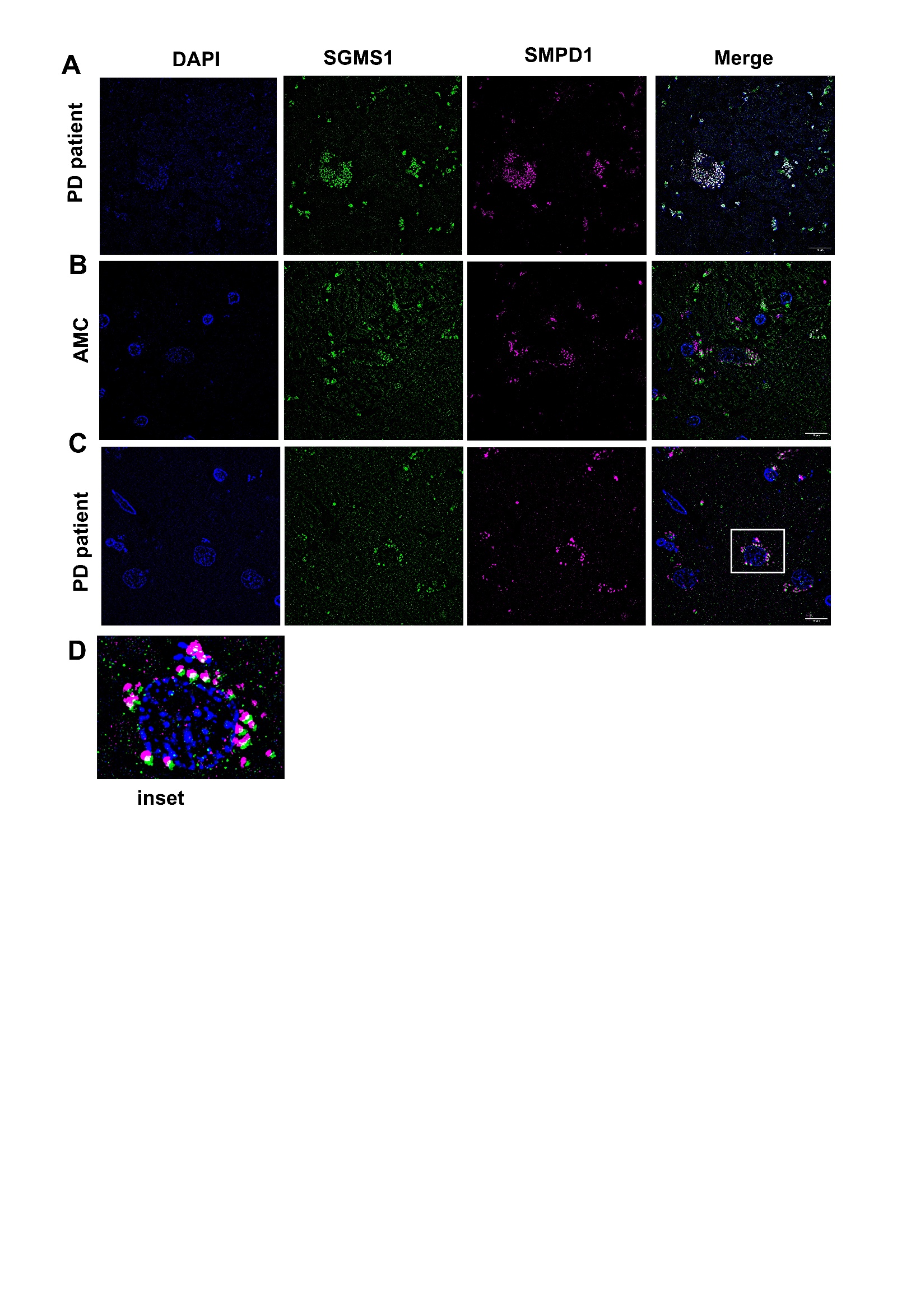

**Figure S8:** Immunofluorescence images for PD patient (A) and Age-matched control (AMC) stained for SMPD1 and SGMS1 (B). Another field for PD patient, showing distinct vacuoles of SGMS1 and SMPD1 (inset, D). Scale bar= 10µm.
